# Antigenic landscape of rabies and related lyssaviruses revealed by cryo-EM

**DOI:** 10.64898/2026.01.14.699513

**Authors:** Heather M. Callaway, Dawid S. Zyla, Kathryn M. Hastie, Stephanie S. Harkins, Shanika Kothalawalage, Nimasha Samarasinghe, Alexander Flynn, Chitra Hariharan, Jieyun Yin, Davide Corti, Herve Bourhy, Scott K. Dessain, Erica Ollmann Saphire

## Abstract

Rabies continues to kill over 60,000 people per year despite life-saving vaccines and post-exposure treatments, and costs billions of dollars in prevention and treatment. Preventing rabies deaths and reducing the global economic burden of the virus will require both developing a monoclonal antibody cocktail to replace human serum in treatment and improving rabies vaccines to elicit long-lasting protection. Here, we solve nine cryo-EM structures of neutralizing monoclonal antibodies in complex with the rabies virus surface glycoprotein (RABV-G). The nine structures span three known antigenic sites plus two new antigenic sites, not previously mapped. We further find that these two new sites are the targets of antibodies with the desired broad neutralization of rabies as well as other emerging lyssaviruses. Across the mAb panel, fusion inhibition and binding affinity correlate best with neutralization. Together, these results provide a roadmap for structure-guided vaccine and therapeutic antibody design for rabies and related lyssaviruses.

## Introduction

Rabies is the most lethal human disease discovered, with a nearly 100% fatality rate for untreated infections. These infections cause over 59,000 deaths annually, with approximately 40% of deaths occurring in children (1). While lifesaving rabies vaccines and post-exposure treatments are available, both have flaws that prevent their universal use. Rabies vaccines elicit only a short-term protective antibody response, with re-vaccination required as frequently as every six months to maintain protective antibody titers(2, 3). As a result, most potential human rabies transmissions are prevented through post-exposure treatment instead of pre-exposure vaccination. Rabies post-exposure treatment consists of a 4- or 5-dose vaccine series administered in combination with human rabies immunoglobulin (HRIG), polyclonal serum derived from the blood of vaccinated human volunteers. Because HRIG is human-derived, it is variable in potency, poses a risk for the transmission of bloodborne pathogens, and can be prohibitively expensive, contributing to fatality rates in the low-income countries where rabies deaths are highest. Both rabies vaccines and HRIG are also specific to rabies virus, showing little or no cross-neutralization for the sixteen rabies-related emerging lyssaviruses, many of which can be transmitted to humans and cause identical clinical symptoms and disease (3–8). To control rabies transmission and develop therapies that are also successful against emerging lyssaviruses, we must develop more durable, broadly neutralizing, and affordable rabies/lyssavirus vaccines and antibody therapeutics. In this work, cryo-EM and functional characterization of nine RABV-G/antibody complexes yield an immunogenic landscape of rabies virus recognition and key information about successful antibodies and epitopes to guide vaccine and therapeutic design.

For multiple viruses, administration of highly purified and well-characterized potent monoclonal antibodies (mAbs) provides a superior treatment option to polyclonal human sera (9–11). Individual mAbs with greater breadth and potency of neutralization than polyclonal serum have been identified for rabies and multiple related lyssaviruses (3, 12). However, these studies lack key structural information about what epitopes antibodies target, what residues they interact with, and how antibody interactions neutralize virus. Having an available epitope map and antigenic landscape of the virus will facilitate selection of antibodies that target spatially distinct regions of the glycoprotein and that avoid regions of high residue variability or frequent escape mutations. High-resolution structural information will also allow directed mutagenesis of antibodies to increase binding strength, neutralization potency, and ability to recognize and neutralize related lyssaviruses.

Rabies virus glycoprotein (RABV-G) is the target of the rabies neutralizing antibody response and consists of three domains: a central domain (CD), a fusion domain (FD), and a Pleckstrin homology domain (PHD) (**Fig. 1**). The central domain contains the α-helix at the core of the glycoprotein and loops which form a trimeric interface with adjacent protomers. The fusion domain contains the fusion loops, and the Pleckstrin homology domain connects the central domain to the fusion domain. On the viral surface, RABV-G adopts multiple conformations (13–17), increasing the number of potential targets for vaccines and therapeutic antibodies. RABV-G reversibly transitions between monomeric, trimeric, pre-fusion, and post-fusion conformations (13–15). During the pre- to post-fusion transition, the central α-helices elongate and the Pleckstrin homology domain rearranges to sit directly above the central domain, changing the availability of some antigenic sites (18, 19). While acidic pH drives this change, the transition is incomplete: some copies of RABV-G remain in the pre-fusion conformation at acidic pH, and some copies maintain the post-fusion conformation when pH is raised to neutral or basic (13). As a result, immunization may result in an antibody response that targets both pre- and post-fusion RABV-G.

**Figure 1.**
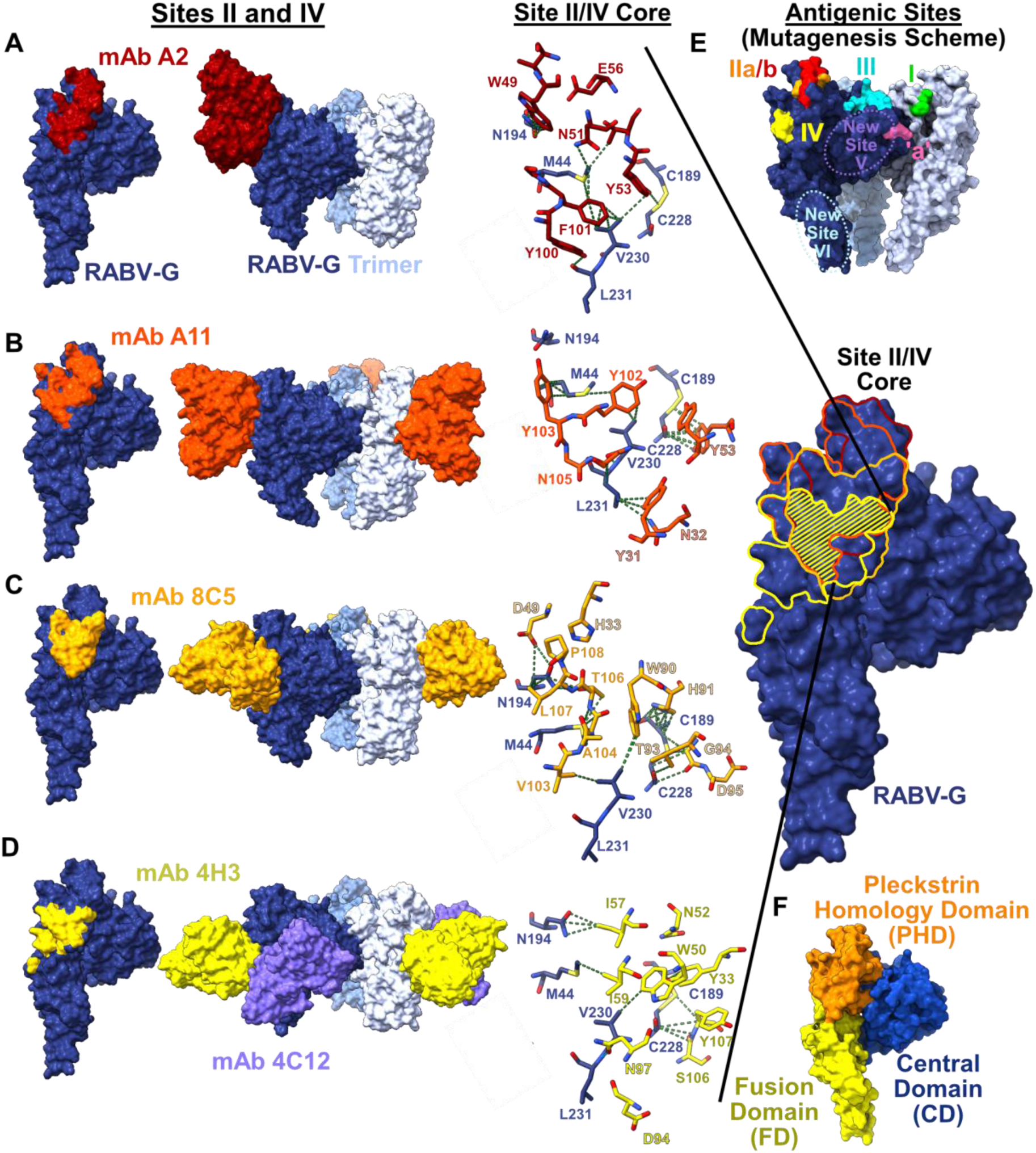
Antibodies at antigenic sites II and IV recognize an overlapping core of amino acids. Mouse mAbs A2 (A) and A11 (B) and human mAbs 8C5 (C) and 4H3 (D) recognize antigenic sites II and IV. Antibody binding footprints (left) are shown depicting all residues on RABV-G within 4Å of the bound antibody. Atomic models of antibodies bound to RABV-G are shown in the middle, and the core region of shared amino acids between all four antibodies is shown to the right. Van der Waals interactions between RABV-G and antibodies are depicted with a green dotted line. Antibody light chain residues are labeled with a lighter shade of red, orange, or yellow and outlined in black. Antigenic sites according to the mutagenesis scheme are shown in (E), with our newly discovered antigenic sites also depicted. Rabies antigenic sites identified by the mutagenesis scheme include residues 263-264 (site I, green); residues 198-200 (site IIa, orange); residues 34-42 (site IIb, red); residues 330-338 (site III, teal); residues 226-231 (site IV, yellow); and residues 342-343 (site ‘a’, pink). RABV-G domains are depicted in (F), with the Pleckstrin Homology Domain (PHD) colored orange, the Central Domain (CD) colored blue, and the Fusion Domain (FD) colored yellow.

Previous work identified several RABV-G antigenic sites, classified either by antibody sensitivity to RABV-G point mutations or by antibody competition. The mutation-based naming scheme identified five major and one minor antigenic site: site I (residues 263-264), site IIa (198-200), site IIb (34-42), site III (330-338), site IV (226-231), and site ‘a’ (342-343) (Fig. 1) (20). The antibody competition-based naming scheme, which used a large antibody panel to define competition groups, separated antibodies as groups I, III, III.2, A, B and C (3). Note that groups named I and III in the competition-based scheme are distinct from antigenic sites I and III in the mutation-based scheme. Both the mutation-based and competition-based naming schemes were developed prior to the availability of 3D structures for RABV-G. Crystal structures of alternate conformation pre- and post-fusion RABV-G became available in 2020 (18), a Pleckstrin Homology Domain/antibody complex in 2020 (21), and the first structures of pre-fusion, trimeric RABV-G in complex with stabilizing antibodies in 2022 (19, 22). Very few antibodies have thus far been illuminated structurally, and therefore many antibody epitopes were largely inferred, rather than directly visualized.

Here, we reveal cryo-EM structures for nine additional mAbs: six at high-resolution (<3.5 Å) and three at intermediate resolution (4.5-7 Å). In this work, the number of new antibodies, combined with, and in comparison to the handful of prior structures, now allow assignment of actual, 3D antigenic footprints to the previously proposed antigenic sites. These nine structures span five antigenic sites, reconcile site nomenclature across the mutagenesis-based and antibody competition-based studies, and include two new antigenic sites not previously identified. Through complementary functional studies, we also identify the likely mechanisms of neutralization against each antigenic site and determine which rabies antibodies recognize glycoproteins from other lyssaviruses. Combining these novel structures and earlier work yields a more complete map of the rabies antigenic landscape, through which we may understand potent and broad rabies and lyssavirus neutralization, identify better candidates for therapeutic antibody cocktails, and select better metrics to evaluate next-generation vaccines for rabies and related lyssaviruses.

## Results

Previously determined structures of monoclonal antibodies (mAbs) in complex with RABV-G include three antibodies that recognize antigenic site III (as defined in the mutation scheme; mAbs RVA122 (19), 17C7 (22), and CTB012 (23)), two that recognizes antigenic site II (RVC20 (21) and 1112-1 (22)), and two that recognize site I (523-11 (18), and CTB011 (23)). In this study, we aimed to characterize additional mAbs to determine the span of neutralizing rabies antibody recognition and identify new rabies antigenic sites.

### MAb discovery and characterization

We synthesized human mAbs from publicly available sequence information (mAbs 4C12, 4H3, 7E8, 8C5, 10H5, and RVC68 (24, 25)) and complemented them with three additional novel mAbs elicited for this study through mouse immunization (mAbs A2, A4, and A11). We then used bio-layer interferometry to organize antibodies into competition groups, observing four groups across the panel: one competition group with mAbs 10H5 and A4 (**Fig. S1**); a second with mAbs 4H3, 7E8, 8C5, A2, and A11; and two more with mAbs RVC68 and 4C12 each forming their own distinct competition groups. Interestingly, 4C12 exhibited directional competition: when placed on the sensor first, it blocks binding of 10H5 and A4, but placing 10H5 and A4 on the biosensor first does not block binding of 4C12. Each of these mAbs were mapped in complex with RABV-G by cryo-EM.

In these studies, we used a wild-type RABV-G which rapidly transitions into conformationally heterogeneous monomers if not stabilized by an antibody. To retain the trimeric, pre-fusion conformation of RABV-G, we employed one of several pre-fusion stabilizing mAbs for structure determination. Binding of mAb RVA122, known to bind site III (as defined by the mutagenic scheme) from our prior work (19), allowed high-resolution structure determination of six additional mAbs against other sites. Our EM studies revealed that 10H5 and A4 also bound site III but did not stabilize trimers as well as mAb RVA122, only allowing intermediate (4.5-7 Å) resolution structures. Even the intermediate-resolution structures, however, allow mapping of the general antibody binding footprints. Overall, nine complexes were determined by cryo-EM, each containing a site III mAb plus one other mAb.

### Rabies antibody binding and new antigenic sites

The previous mapping by mutagenesis, before the availability of RABV-G structures, identified residues on RABV-G important for antibody binding and clustered them into four antigenic sites. Residues 263-264 were previously named “antigenic site I”, residues 34-42 and 198-200 “antigenic site II”, residues 330-338 “antigenic site III”, and residues 226-231 “antigenic site IV” (Fig. 1) (20). Structures of RABV-G in complex with RVC20 (site 1 in the competition-based scheme) (21) and RVA122 (site III in the competition scheme) (19) have linked competition group I to the Pleckstrin-homology domain (sites II and IV from the mutagenesis-based naming scheme) and competition group III to site III from the mutagenesis-based scheme. The nine cryo-EM complexes described here substantially expand on that work. Five antibodies from these nine complexes target antigenic sites II and IV, revealing an overlapping epitope rather than two distinct antigenic sites. Two other mAbs recognize epitopes containing residues associated with antigenic site III. The remaining two antibodies recognize new epitopes not overlapping with any classical antigenic site, which we have named ‘antigenic site V’ and ‘antigenic site VI’, expanding on the classic mutagenesis-based nomenclature.

**Antigenic sites II and IV:** Human antibodies 8C5 (3.16 Å), 4H3 (3.24 Å), and 7E8 (∼4.5 Å), and murine antibodies A2 (3.01 Å) and A11 (2.92 Å) recognize the Pleckstrin homology domain (PHD), with footprints spanning residues associated with antigenic sites II and IV (**Fig. 1**). Although these residue clusters were previously identified as distinct sites via mutagenesis (20), these five structures show that antibodies recognize these residues not in two distinct groups, but instead in an overlapping ∼25 nm^2^ continuum, with mAb A2 at the apex and mAb 4H3 toward the base (Fig. 1). We built atomic models of the four site II/IV antibodies that reached high resolution (Fig. 1) to identify the shared amino acid contacts. In these models, we observed that all four site II/IV antibodies with high-resolution structures form Van der Waals interactions with a shared set of contacts on RABV-G including residues M44, C228, and V230. Details and additional contacts, including van der Waals interactions of antibodies A2, A11, 8C5, and 4H3 with C189, N194, and/or L231, are illustrated in Figure 1. Each of these site II/IV mAbs binds both pre- and post-fusion RABV-G (**Fig. S2**); the Pleckstrin homology domain that they target moves as a rigid body in the conformational transition(18, 19).

**Antigenic site III:** Human mAb 10H5 (6.69 Å) and murine antibody A4 (∼4.5 Å) both recognize the central domain (CD) of RABV-G with footprints that overlap residues associated with antigenic site III (mutagenic scheme) and the RVA122 epitope (19, 20) (**Fig. 2**). All three of these antibodies recognize a conformational epitope only present on the pre-fusion conformation. Although both 10H5 and A4 only bind pre-fusion RABV-G and we observed pre-fusion trimers during imaging, there was greater structural heterogeneity in A4/RABV-G and 10H5/RABV-G complexes than in complexes made with RVA122 (**Fig. S3-8**). As a result of this and lower number of pre-fusion RABV-G trimers **(Table S1)**, neither 10H5 nor A4 reached a resolution high enough to build an atomic model. The intermediate-resolution footprints, however, suggest that 10H5 and A4 bind solely to the central domain, while in contrast, RVA122 recognizes an epitope bridging the central and Pleckstrin homology domains, likely better stabilizing the pre-fusion trimer.

**Figure 2.**
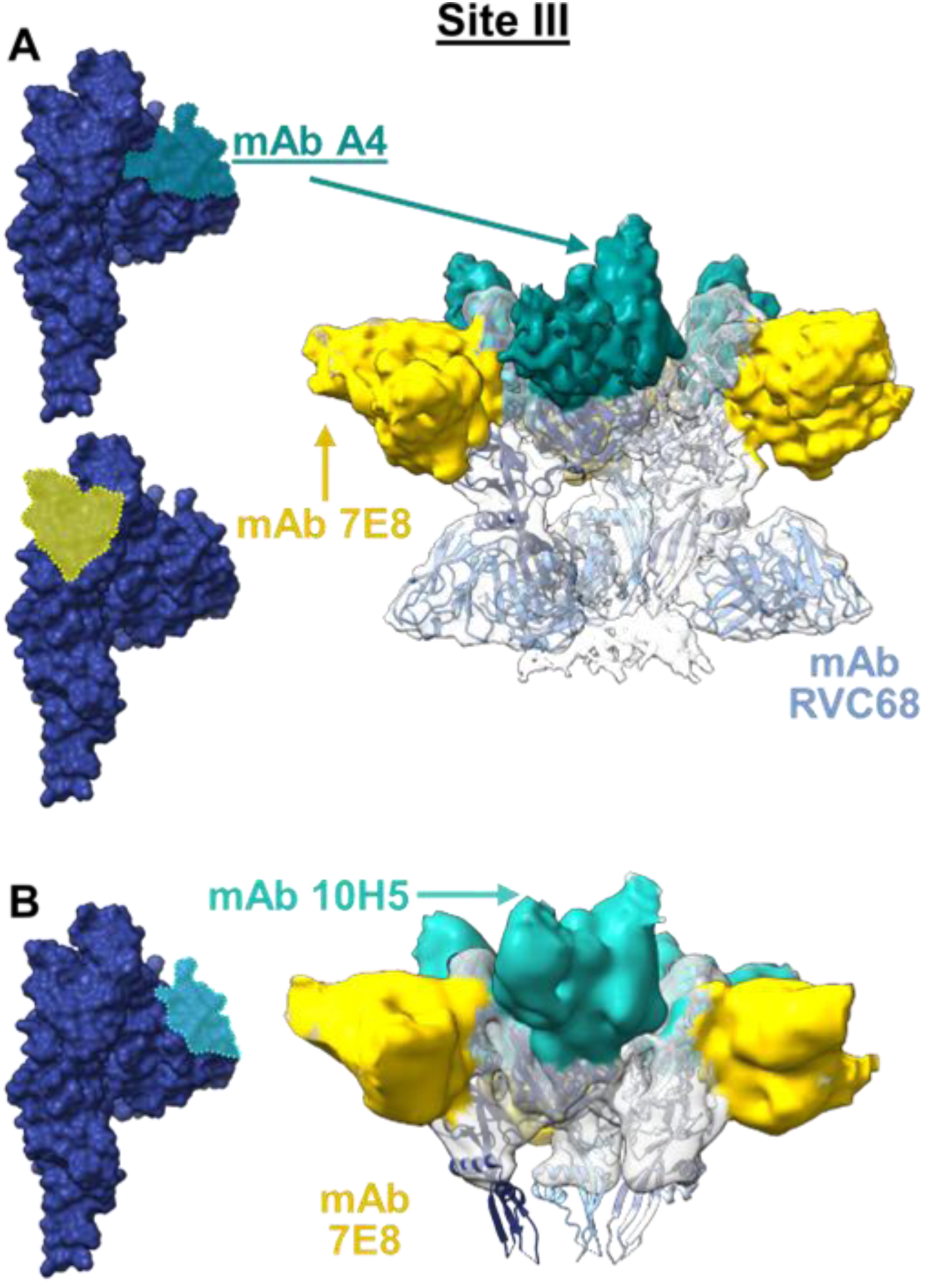
Site III monoclonal antibodies stabilize the pre-fusion conformation of RABV-G. Low resolution cryo-EM maps of site III mAbs A4 (mouse) (A) and 10H5 (human) (B) show that site III-binding mAbs other than RVA122 also recognize the pre-fusion conformation of RABV-G. Approximate binding footprints for each mAb are shown on the left. Intermediate-resolution cryo-EM maps (right) are shown with new mAbs colored teal or yellow, and RABV-G and mAb RVC68 models docked into the map. An additional site II/IV human mAb, 7E8 (yellow), and its binding footprint are also shown. 7E8 is not depicted in the site II/IV Figure 1 due to its lower resolution.

**New antigenic site V:** Human mAb 4C12 is also specific for the pre-fusion conformation of RABV-G, but does not compete with RVA122 (Fig. S6). The cryo-EM structure of mAb 4C12 (3.24 Å) revealed a new antigenic site below antigenic site III, not associated with any residues previously identified in RABV-G mutagenesis studies. (**Fig. 3a**). We term this antigenic site ‘site V’ as a continuation of the classical rabies antigenic site nomenclature (**Fig. 3a**). Here, the 4C12 footprint bridges the lower central domain (CD) to residues W12, P13, P16, and N57 in the Pleckstrin homology domain (PHD). This anchoring of the CD and PHD is similar to the pre-fusion trimer stabilizing effect we previously observed with RVA122 (19). Notably, 4C12 is the only pre-fusion-specific antibody yet characterized that does not recognize antigenic site III.

**Figure 3.**
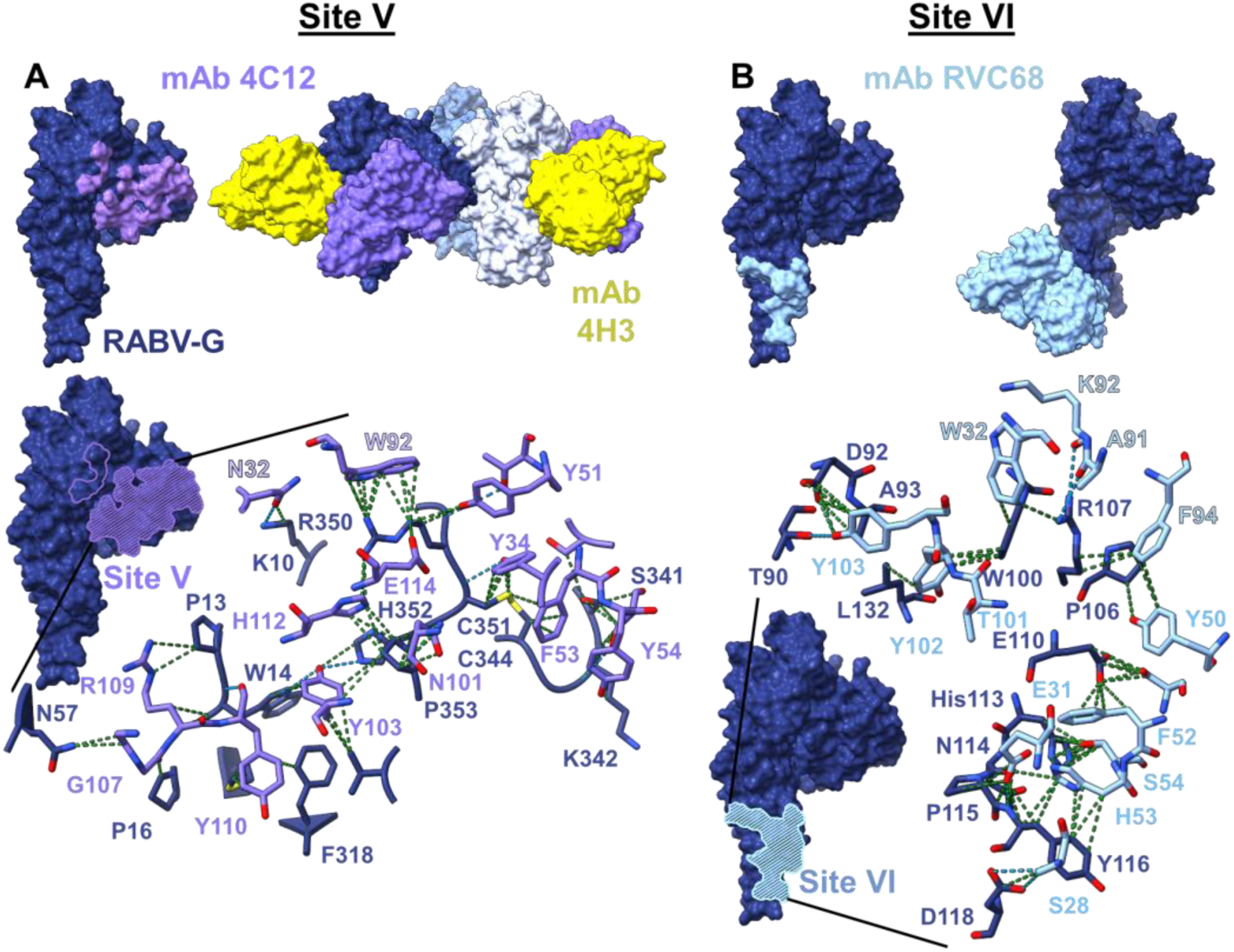
New antigenic sites on the central domain (site V) and fusion domain (site VI). Human mAbs 4C12 (A) and RVC68 (B) recognize new binding sites on the central domain and fusion domain, respectively. Antibody binding footprints (top left) are shown depicting all residues on RABV-G within 4Å of the bound antibody and atomic models of RABV-G/antibody complexes are shown on the top right. A magnified view of RABV-G/antibody binding interactions (bottom) is shown for each antibody, with Van der Waals interactions depicted with a green dotted line and hydrogen bonds with a light blue dotted line. Glycoprotein residues are depicted in dark blue, and antibody residues in purple (site V) or cyan (site VI). Antibody light chain residues are labeled with a lighter shade of purple or blue and outlined in black.

**New antigenic site VI:** mAb RVC68 does not compete with any other mAb in the study. The cryo-EM structure of its complex reveals another new antigenic site, we term ‘site VI’, on the fusion domain (FD) (**Fig. 3b**). RVC68 recognizes RABV-G via four hydrogen bonds and an extensive network of Van der Waals interactions, with almost half of the interactions contributed by six aromatic amino acids in the RVC68 paratope. Residues W32 in CDRL1, F94 in CDRL3, Y50 and F52 in CDRH2, Y102 and Y103 in CDRH3 mediate contact to RABV-G residues 110-118 and the horizontal α-helix adjacent to them (Fig. 3b). RVC68 binds RABV-G at both neutral and acidic pH (pre- and post-fusion conformations, respectively), with more binding at acidic pH (Fig. S2). While the residues in the RVC68 epitope maintain the same shape in both the pre-and post-fusion conformations of RABV-G(18, 19), the post-fusion conformation may be more sterically accessible to IgG.

### Broad Reactivity

A major goal of rational rabies therapeutic antibody and vaccine design is to develop therapeutics effective not only against rabies, but also the many rabies-related lyssaviruses that cause the same clinical disease as rabies virus. To determine whether specific antigenic sites or amino acid contacts facilitate broad recognition of rabies-related lyssaviruses, we screened our panel of mAbs for binding to glycoproteins from seven additional lyssaviruses. We chose glycoproteins (G) from lyssaviruses across all four lyssavirus phylogroups to measure broad antibody recognition. These included four members of phylogroup I, the largest group which also includes rabies virus [Duvenhage lyssavirus (DUVV), Eastern Bat Lyssavirus-1 (EBLV-1), Australian Bat Lyssavirus (ABLV) and Irkut Lyssavirus (Irkut)]; one member of phylogroup II [Mokola lyssavirus (Mokola)]; one member of phylogroup III [West Caucasian Bat Lyssavirus-1 (WCBLV-1)]; and one member of phylogroup IV [Ikoma lyssavirus (Ikoma)].

Because several of these glycoproteins do not produce sufficiently infectious pseudoviruses for neutralization assays in our hands or express well as soluble glycoprotein ectodomains, we assayed antibody binding against full-length glycoproteins expressed on 293T cells (**Fig. 4**). To ensure that glycoproteins reached the cell surface and folded correctly, we tested them in a fusion assay with a green fluorescent reporter as the output for successful fusion (**Fig. 4, right**). Two glycoproteins, Eastern Bat Lyssavirus 1-G and Mokola-G, failed to induce cell-cell fusion in this assay, but were both recognized by antibodies against conformational and quaternary epitopes (RVC68 against both glycoproteins and two different group III mAbs – 4C12 for Eastern Bat Lyssavirus-1-G and 10H5 for Mokola-G), indicating that these glycoproteins did reach the cell surface in a properly folded form.

**Figure 4.**
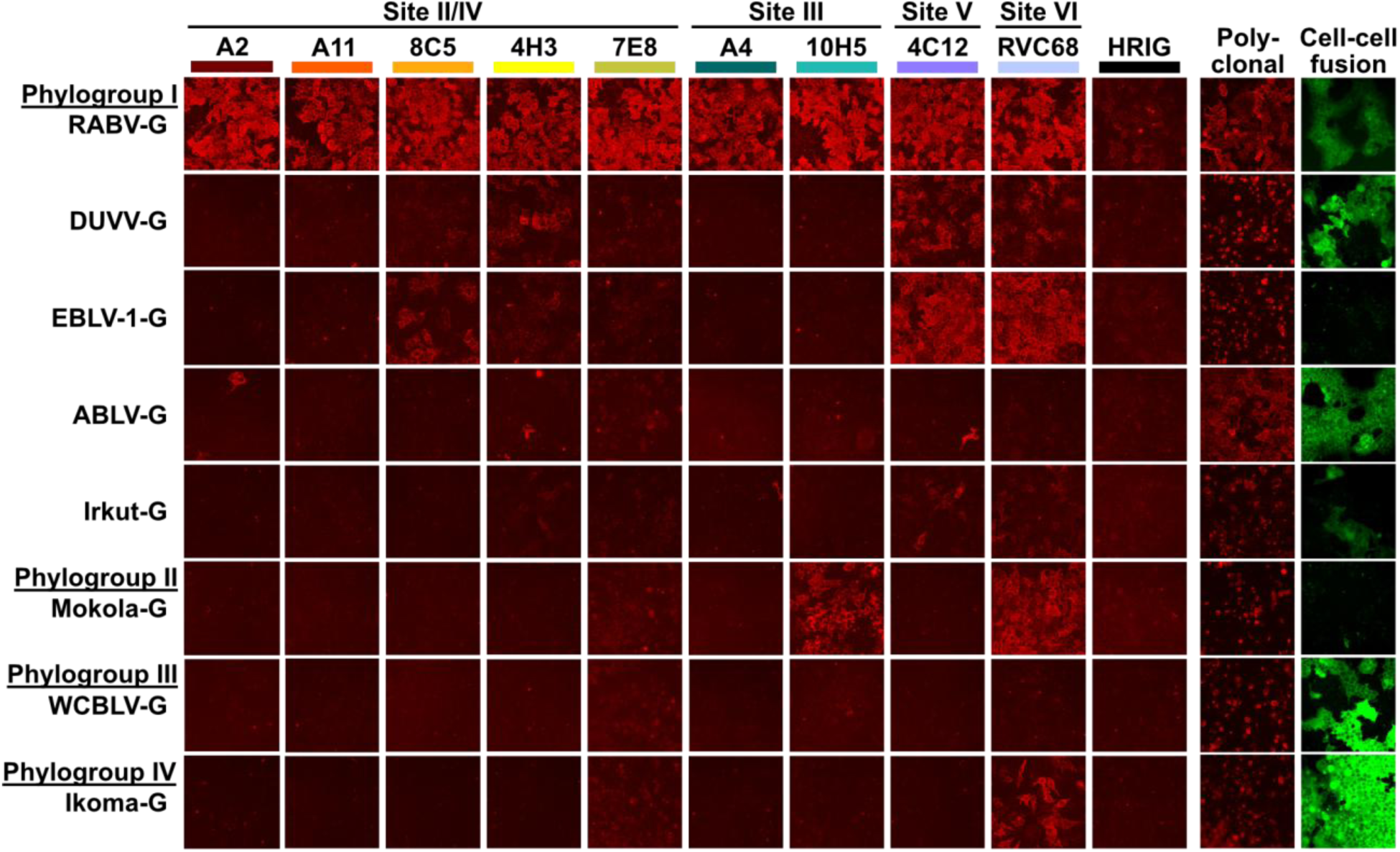
Site V and VI rabies monoclonal antibodies broadly recognize other lyssavirus glycoproteins, while Site II/IV and III are largely rabies-specific. Binding of mAbs to full-length lyssavirus glycoproteins expressed on 293T cells indicate that some rabies antibodies bind broadly. A polyclonal anti-rabies antibody and a cell-cell fusion assay (right) confirmed that each full-length glycoprotein expressed and reached the cell surface. While not all glycoproteins induced cell-cell fusion in our assay, the two that did not (Eastern Bat Lyssavirus-1 G and Mokola-G) were recognized by mAbs, indicating that they reached the cell surface and were well-folded.

Of the antibodies in our panel, site VI antibody RVC68 was by far the most broadly cross-reactive, in keeping with previous reports (3). In our assay, RVC68 bound to glycoproteins from Duvenhage, Eastern Bat Lyssavirus-1, Irkut, Mokola, and Ikoma lyssaviruses (Fig. 4). The broad cross-reactivity we and others have observed for RVC68 almost certainly results from the broad sequence conservation of its epitope across lyssaviruses (**Fig. 5**). The only glycoproteins that RVC68 did not recognize in our assay were Australian Bat Lyssavirus-G and West Caucasian Bat Lyssavirus-G. Previous reports indicate that Australian Bat Lyssavirus neutralization was strain specific (3), and while the RVC68 contacts are conserved in Australian Bat Lyssavirus-G, a lysine residue in the footprint at position 96 appears likely to disrupt binding. For West Caucasian Bat Lyssavirus-G, in contrast, the RVC68 binding site is much less conserved. Based on sequence comparison, the most disruptive non-conserved residue is likely at W100 (R in West Caucasian Bat Lyssavirus-G), which has also been shown to disrupt rabies neutralization in a deep mutational scanning assay (26).

**Figure 5.**
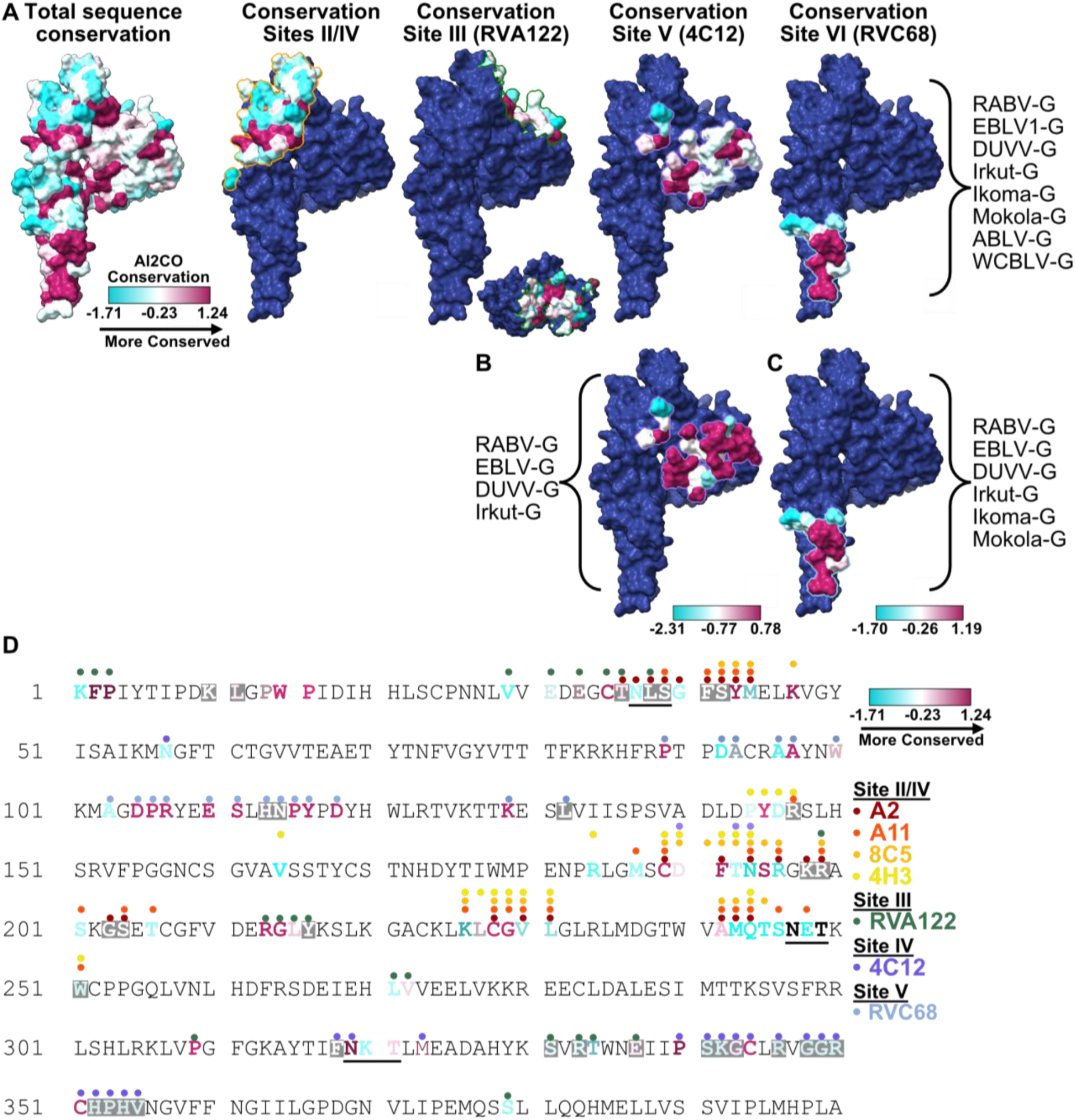
Rabies antibody cross-reactivity likely results from sequence conservation in the antibody binding site. Amino acid sequence conservation across eight lyssavirus glycoproteins was projected onto RABV-G (A), with binding footprints for antibodies at each antigenic site outlined and site-specific sequence conservation shown. Site III recognition is to mAb RVA122 as the other site III mAbs are too low in resolution to map specific contacts. For antibodies 4C12 at antigenic site V (B) and RVC68 at antigenic site VI (C), sequence conservation was shown among cross-reactive lyssavirus glycoproteins. Sequence conservation across RABV-G antibody binding sites was also depicted on the amino acid sequence (D), with residues within 4Å of antibodies shown with color-coded circles. Amino acid sequence conservation for each contact is also shown for RABV-G, and glycosylation sites are underlined.

The second most cross-reactive antibody in our panel, site V antibody 4C12, recognized glycoproteins from Eastern bat lyssavirus 1, Duvenhage, and Irkut lyssaviruses (Fig. 4). The 4C12 binding footprint is largely conserved among rabies and these lyssaviruses (Fig. 5b), with only two residues that interact with 4C12 substantially differing from rabies (residues 318 and 355, at the edge of the binding footprint). We observed similar conservation of binding sites for site II/IV antibodies 8C5 and 4H3, which recognize glycoproteins from Eastern bat lyssavirus 1 and Duvenhage viruses, respectively. Of the glycoprotein residues interacting with 8C5, only two substantially differ between RABV-G and Eastern Bat Lyssavirus 1-G (residues 194 and 242; N and A in RABV-G, and T and S in Eastern Bat Lyssavirus 1-G). A larger number of differences between RABV-G and Duvenhage-G exist in the 4H3 binding footprint, but only one residue that interacts with 4H3 substantially differs between the glycoproteins (residue 194; N in RABV-G and R in Duvenhage-G).

Site III antibody 10H5, finally, is notable because only 10H5 and RVC68 recognize Mokola glycoprotein. While we hypothesize that sequence similarity between RABV-G and Mokola-G in the antibody binding site drives cross-reactivity, our current structure lacks the resolution to identify specific amino acid contacts between the antibody and the glycoprotein.

Overall, we find that the two antibodies at new antigenic sites V and VI have the broadest cross-reactivity, whereas antibodies recognizing sites II/IV and III are largely rabies-specific. Because the residues present in sites V and VI are well conserved among lyssaviruses, we hypothesize that other antibodies that bind these sites will show similar cross-reactivity.

### Potency and Mechanisms of Neutralization

In addition to determining structures of RABV-G/antibody complexes, we also measured antibody neutralization potency and determined mechanisms of neutralization. By comparing antibodies within and between antigenic sites, we sought to determine if antibodies that recognized specific sites (or residues within those sites) neutralized virus with the same strength and mechanism, and if any antigenic sites elicit more potent antibodies.

As IgGs, all the antibodies in our panel except for RVC68 neutralized rabies pseudoviruses potently, with IC50s of approximately 10pM (**Fig. 6a**). Broadly reactive site VI mAb RVC68, in contrast, was 100-fold less potent than the other eight antibodies. IgG and Fab neutralization was equivalent for antibodies 7E8 (site II/IV), 10H5 (site III), and RVC68 (site VI), suggesting that these antibodies function by binding glycoprotein monovalently. For other antibodies, neutralization was inferior in the Fab format, indicating that for these antibodies, antibody bivalency and glycoprotein crosslinking enhances neutralization (**Fig. 6b**). This finding occurred for antibodies A2, A4, A11, 4C12 and 4H3 in both antigenic groups II/IV and V. The most potent of the Fabs was the site III binder 10H5, which neutralized just as well as its IgG form, followed by site II/IV binders 8C5 and 4H3 and site V binder 4C12 (Fig. 6a). Notably, Fab binding affinity to RABV-G (**Fig. S1**) did correlate with Fab neutralization potency, with antibodies with the highest affinity neutralizing the most potently and antibodies with the lowest affinity neutralizing least potently.

**Figure 6.**
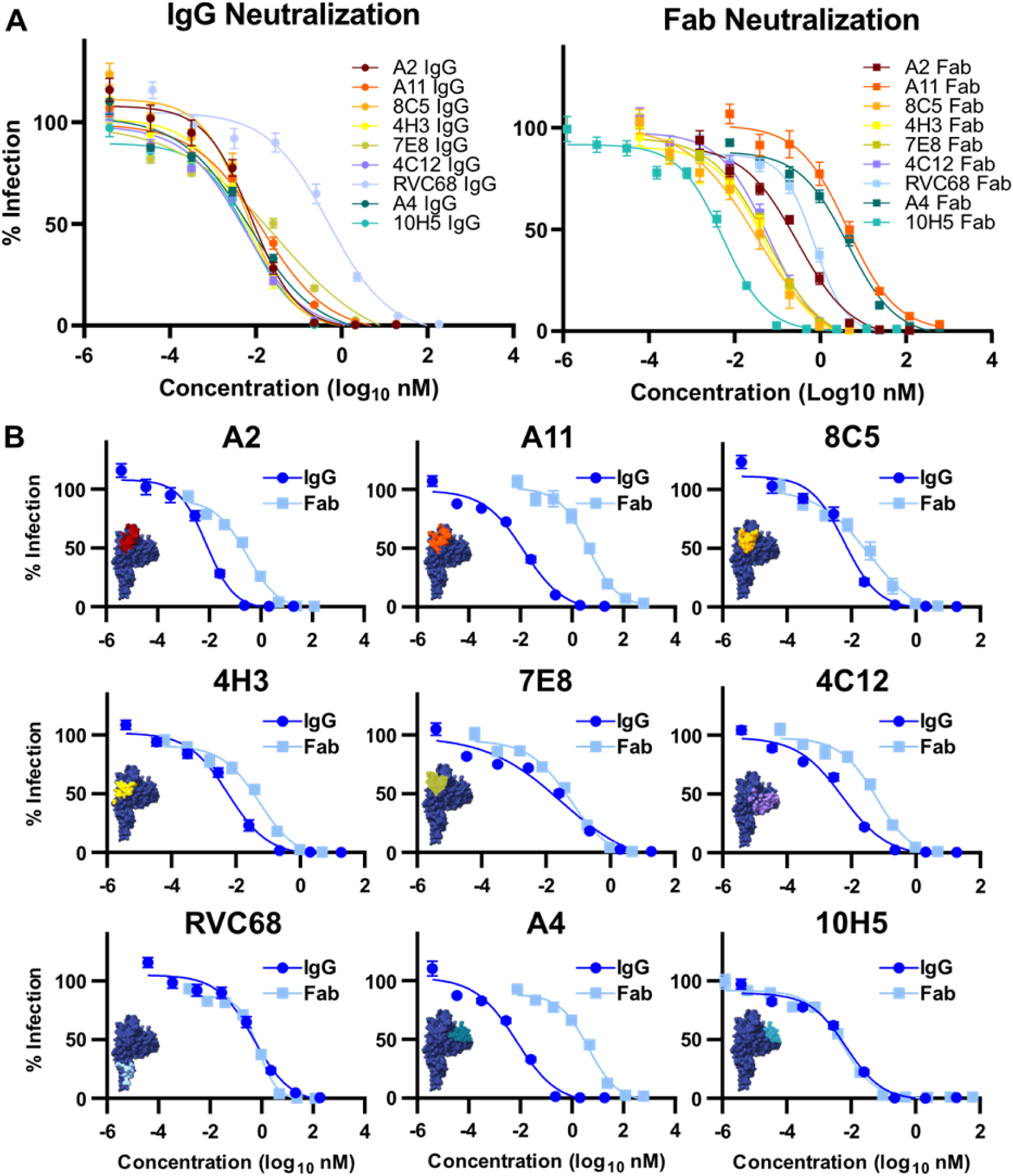
For most rabies antibodies, IgG bivalency enhances neutralization potency. IgG and Fab neutralization of rabies pseudoviruses is shown comparing all IgGs and Fabs (A), and for individual monoclonal antibodies (B). A substantial difference between IgG and Fab neutralization indicates that antibody bivalency enhances neutralization. The requirement for IgG bivalency in neutralization is not specific to any antigenic site.

As many of these antibodies neutralize better as IgG, we sought to determine which epitopes might be capable of inter- or intra-spike recognition. We aligned structures of whole IgGs from the protein databank with our models, but only antibody RVA122 from our original rabies antibody binding study (19) appears to bind in a way that may allow intra-spike binding to the RABV-G pre-fusion trimer. While a structure of the RABV-G post-fusion trimer has not yet been determined, we do not expect that any of these antibodies can bind it with bivalent, intra-spike recognition. The site II/IV antibodies would not be able to reach a second protomer in a trimer, our site III and V antibodies are pre-fusion specific, and the site VI antibody RVC68 does not neutralize better as an IgG. Instead, it is likely that antibodies that neutralize better as IgGs than Fabs are capable of inter-spike recognition.

We next performed binding and uptake assays with IgGs to determine if antibody binding blocked pseudovirus uptake into 293T cells. Both site III antibodies A4 and 10H5 and the sole site VI antibody RVC68 block binding and uptake. The sole site V antibody, 4C12, does not block binding and uptake. The site II/IV antibodies were split: 7E8 successfully blocks binding and uptake while A2, A11, 8C5, and 4H3 do not (**Fig. 7**). In addition, every antibody that neutralized equally well as a Fab and IgG (Fig. 6b) blocked binding and uptake.

**Figure 7.**
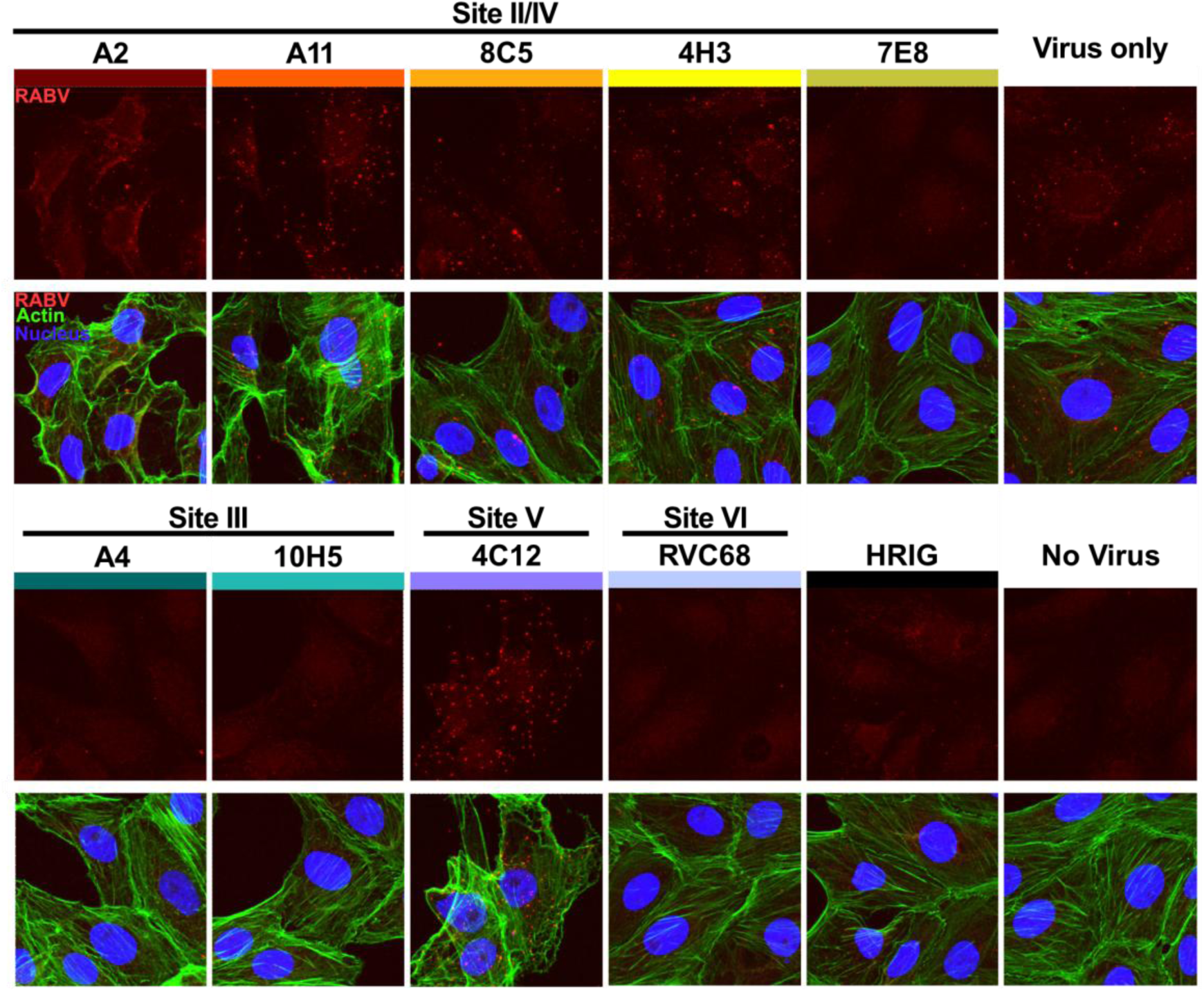
Site III, V, and VI monoclonal antibodies block uptake into 293T cells; except for mAb 7E8, site II/IV antibodies do not. Depicted are binding/uptake assays for site III mAbs A4 and 10H5; site V mAb 4C12, and site VI mAb RVC68 which block binding and uptake; as well as site II/IV mAbs A2, A11, 8C5, 4H3 which do not block binding and uptake. 7E8 is the only site I/IV mAb that blocks binding and uptake of virus. Binding and uptake assay of rabies pseudoviruses with or without monoclonal antibodies into 293T cells. Pseudoviruses are stained red, actin (via phalloidin) is stained green, and nuclei (Hoescht) is stained blue.

Finally, we used a fusion inhibition assay with surface-displayed RABV-G to determine how well IgGs and Fabs blocked viral fusion. We found that all IgGs, including those that neutralized equally well as Fabs and IgGs, potently inhibited RABV-G fusion (**Fig. 8**). The steric bulk of the IgG itself may assist in inhibiting conformational changes required for membrane fusion. Fabs, in contrast, varied substantially in how well they inhibited fusion, with no correlation to antibody binding site. Fabs from site II/IV antibodies 8C5, 4H3, and 7E8 inhibited fusion strongly, while other site II/IV antibodies A2 and A11 were much less effective. Similarly, among site III antibodies, 10H5 Fabs were strong fusion inhibitors, while A4 Fabs barely blocked fusion at all. Antibody binding at low pH (**Fig. S2**) also showed no correlation with Fab fusion inhibition. Overall, those Fabs that neutralize well also block fusion well, and those that neutralize poorly also block fusion poorly.

**Figure 8.**
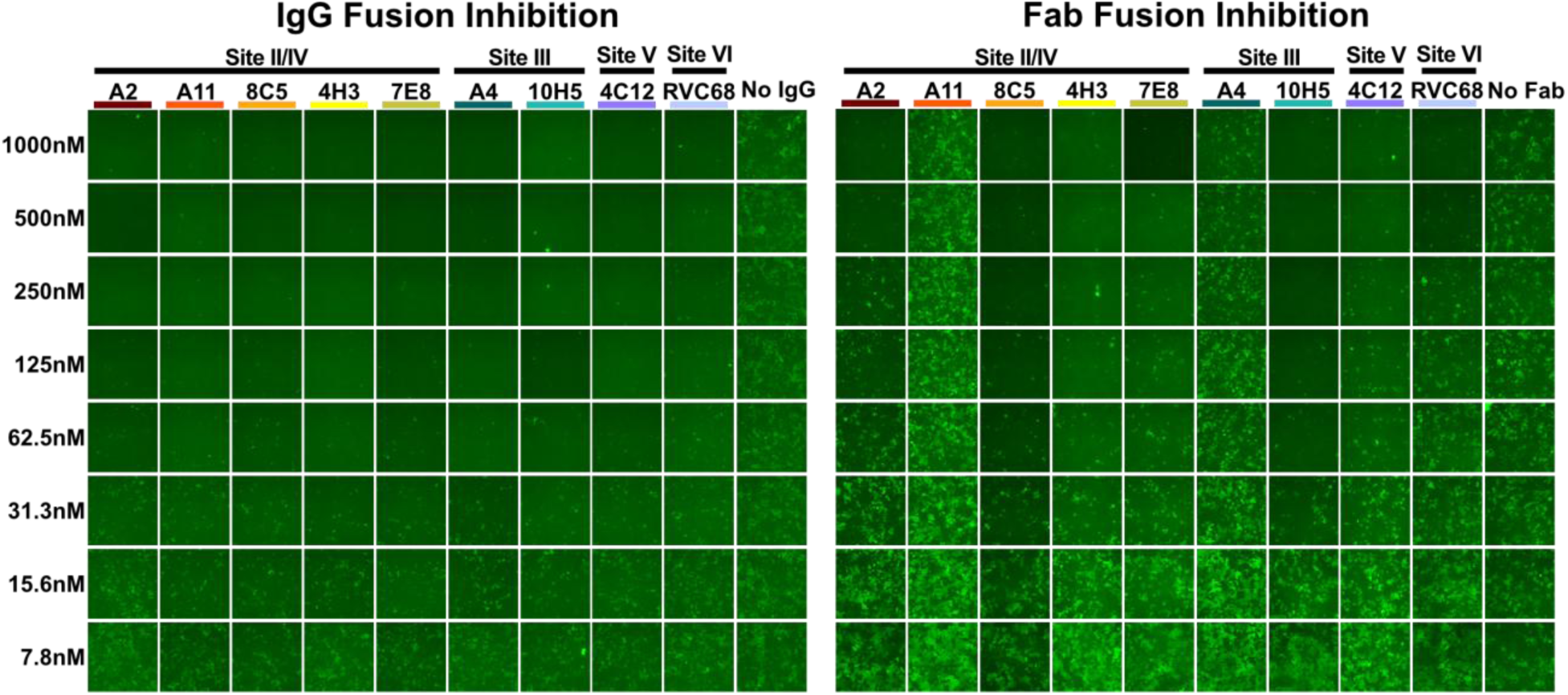
While all rabies IgGs potently inhibit fusion, Fab fragments vary in potency. Fusion inhibition assay using IgGs (left) and Fabs (right). 293T cells expressing half of a split mNeon Green protein were co-seeded into plates, transfected with RABV-G, and then incubated with serially diluted IgGs or Fabs before exposure to acidic pH. Green fluorescence indicates successful fusion of cell membranes.

Together, our results indicate that rabies antibody neutralization potency correlates with binding affinity and fusion inhibition. The Fabs that neutralize most potently (10H5 and 8C5) bind with the highest affinity and inhibit fusion strongly, while the Fabs that are the least potent (A4 and A11) bind with lower affinity and weakly inhibit fusion.

## Discussion

Here, we solved structures of nine new rabies antibodies in complex with RABV-G, expanding the breadth of structural information available for antibody binding at antigenic sites II, III, and IV, and mapping new antigenic sites V and VI. Importantly, we find that the newly mapped sites V and VI are unusually well conserved among lyssaviruses, and may help immunotherapeutic cocktails achieve broad lyssavirus neutralization. Further, the larger number of structures described here reveal that sites II and IV are recognized by a continuum of antibodies rather than discrete clusters of footprints. We also find that fusion inhibition, likely enhanced by binding affinity and IgG avidity, is the strongest correlate of mechanical neutralization. Together, this work maps the rabies antibody binding landscape and serves as a foundation for design of future structure-guided rabies and pan-lyssavirus vaccines and therapeutic antibody cocktails.

Based on the number of antibodies mapped to each antigenic site (3, 19, 21, 22), the immunodominant epitopes on RABV-G are antigenic sites II/IV and III. However, residues contained in sites II and IV are poorly conserved among rabies-related lyssaviruses, and the site III antibodies in this study, 10H5 and A4, proved similarly rabies-specific. While some site II/IV and III binding antibodies have shown broader recognition of phylogroup I lyssaviruses (3), binding is also antibody and strain specific. In contrast to the immunodominant epitopes, new antigenic site V and VI antibodies, although rarer, are substantially more broadly reactive. Site VI antibody RVC68, in particular, recognizes an extremely well-conserved binding site. Of the glycoproteins from our panel, only Australian Bat Lyssavirus glycoprotein and West Caucasian Bat Lyssavirus glycoprotein (the latter likely because of a W100R point mutation (26)) were not recognized. Although not quite as broadly reactive, site V antibody 4C12 recognized three of four phylogroup I lyssaviruses we tested (those more closely related to rabies) and had good sequence conservation across its binding site. While site V and VI antibodies are rarer than those recognizing immunodominant epitopes, their potential for broader neutralization makes them desirable in a vaccine response or therapeutic cocktail.

Fusion inhibition was the strongest determinant of antibody neutralization in this study. However, whether an antibody recognized only pre-fusion RABV-G or both glycoprotein conformations was not a requirement for neutralization. Site III and V monoclonal antibodies recognize epitopes that are only present in the pre-fusion conformation and presumably block fusion by inhibiting the transition to the post-fusion conformation. In contrast, site II/IV and VI antibodies recognize epitopes present in both pre- and post-fusion RABV-G and inhibit fusion even as Fabs. These antibodies bind well at acidic pH, but as their epitopes likely preclude intra-trimer binding for the post-fusion glycoprotein, we suggest that they block fusion either by preventing post-fusion trimers from forming or by crosslinking glycoproteins from different trimers or other assemblies. Notably, some antibodies that recognized both glycoprotein conformations were just as potent as pre-fusion specific antibodies, suggesting that improved rabies vaccines may not require glycoprotein exclusively in the pre-fusion conformation.

A secondary determinant of antibody neutralization potency was the angle of attachment. Within site II/IV, the less potently inhibiting antibodies A2 and A11 bind to the glycoprotein at a 90° angle compared to the better inhibiting antibodies 8C5, 4H3 and 7E8 (Fig. 1 and S4-7). Antibody binding angles that support bivalent IgG binding, strengthening binding through avidity and inhibiting fusion either sterically or through glycoprotein crosslinking, may also compensate for less potently inhibiting Fabs.

We therefore expect that a good strategy to engineer an antibody for improved fusion inhibition and more potent viral neutralization would first require selecting an antibody that is either pre-fusion specific or binds at a site and angle that prevent formation of post-fusion trimers. Point mutations to that antibody could then introduce bonds to increase binding affinity, breadth of neutralization, and the efficacy of escape mutations. The current standard of care in rabies post-exposure antibody prophylaxis is polyclonal human serum (human rabies immunoglobulin - HRIG) to neutralize virus at the wound site. A mAb cocktail against different, complementary epitopes, and composed of members that each have broad specificity, will provide a more potent, better characterized, more reproducible, and more broadly reactive option for treatment than current HRIG. Indeed, for rabies virus, monoclonal antibodies have been observed to be more potent and broadly reactive than polyclonal sera (3). Recent work has proposed a two-antibody cocktail to replace HRIG (27), consisting of RVC20, which binds at antigenic sites II/IV (21) and RVC58 (3), which binds to antigenic site III. Both antibodies have been shown to potently neutralize phylogroup I lyssaviruses (3, 27).

Our work on rabies epitope mapping and mechanisms of antibody neutralization show that there is physical space for a third antibody bridging the central and fusion domains, either at site V or the more conserved site VI, in this and other cocktails. Including a combination of Pleckstrin Homology Domain (PHD), Central Domain (CD) and Fusion Domain (FD)-binding antibodies could allow for greater breadth of neutralization through targeting different sets of conserved residues for the four lyssavirus phylogroups at each domain. For example, a cocktail consisting of a broadly reactive PHD-binding antibody such as RVC20 (3, 21), a broadly reactive FD-binding antibody such as RVC68 and a broadly reactive CD-binding antibody such as RVC58, 10H5, or 4C12 would reduce the likelihood of escape mutations and increase the number of complementary mechanisms of lyssavirus neutralization.

Together, these results set forth the antigenic landscape of RABV-G and form a foundation for the improved rabies and pan-lyssavirus vaccines and rational design of rabies monoclonal antibody cocktails to replace HRIG. The antibody-antigen cryo-EM structures presented here also provide templates for guidance and interpretation of rational or combinatorial improvements of the individual antibodies themselves for use as therapeutic agents, and for vaccine development as well.

## Acknowledgments

We thank a Tullie and Rickey Families SPARK Award for Innovations in Immunology (HMC) and institutional funds of La Jolla Institute for Immunology (EOS) and Montana State University (HMC) for financial support, and Dr. Ruben Diaz and the cryo-EM facility of La Jolla Institute for Immunology (LJI) for data collection. The Zeiss LSM 880 confocal microscope was funded by NIH S10 OD021831. Instrumentation of the LJI cryo-EM facility is supported by U19 AI109762-S1, gifts from GHR Foundation and private philanthropic support.

## Data availability

Cryo-EM density maps were deposited in the Electron Microscopy Data Bank under EMDB accession codes EMD-xxxxx, EMD-xxxxx, EMD-xxxxx, EMD-xxxxx, EMD-xxxxx, EMD-xxxxx, EMD-xxxxx, and EMD-xxxxx, and atomic model coordinates were deposited in the Protein Data Bank under the PDB accession codes yyyy, yyyy, yyyy, yyyy, and yyyy.

## Methods

### Cells and plasmids

Soluble RABV-G ectodomains and pseudoviruses used in neutralization assays were produced in 293T cells (Homo sapiens; American Type Culture Collection, CRL-3216) grown in Dulbecco’s modified Eagle’s medium (DMEM; ThermoFisher Scientific) with 10% fetal bovine serum (FBS) and 1% penicillin/streptomycin (ThermoFisher Scientific) at 37°C and 5% CO_2_.

Pseudovirus titration and neutralization assays were performed in Vero cells (Cercopithecus aethiops; American Type Culture Collection, CCL-81) grown in DMEM with 10% FBS. Whole IgGs were expressed in ExpiCho cells (Cricetulus griseus; ThermoFisher Scientific) grown in ExpiCHO Expression Medium (ThermoFisher Scientific) at 37°C and 8% CO_2_ in a rotary shaker. Fab fragments with C-terminal strep tags were prepared in S2 insect cells (Drosophila melanogaster; Invitrogen, R69007) grown in Insect-XPRESS medium (Lonza) with 1% penicillin-streptomycin (ThermoFisher Scientific) at 27°C in a rotary shaker.

### Full length glycoproteins and strains

Full length, wild-type glycoproteins for rabies and seven additional lyssaviruses were synthesized with human codon optimization under the CMV promoter in the pCDNA 3.1(-) backbone. These glycoproteins included PV-strain RABV-G (PDB: 7u9g), Irkut lyssavirus glycoprotein (Genbank: AFP74571), Ikoma lyssavirus glycoprotein (Genbank: YP_006742183), Eastern bat lyssavirus 1 glycoprotein (Genbank: AAK97857 with L244Q, P520A, and P521T), Duvenhage lyssavirus glycoprotein (Genbank: ACF32424 with S244L, H322R, R498M; numbering includes signal peptide), Australian Bat Lyssavirus (Genbank: NP_478342.1 with E432K) Mokola lyssavirus glycoprotein (Genbank: AAA67271), and West Caucasian bat lyssavirus glycoprotein (Genbank: YP_009094271).

### Antibody production and purification

To express whole IgGs, antibody variable regions were cloned into phCMV expression vectors containing the human IgG1 heavy chain, kappa light chain, or lambda light chain. IgGs were expressed in ExpiCho cells via transfection with ExpiFectamine (ThermoFisher Scientific) and 15µg plasmid DNA (8µg light chain and 7µg heavy chain) in 25mL culture volume according to the manufacturer’s protocol. Antibodies were purified via a Protein A affinity chromatography column (Cytiva) and buffer exchanged into PBS prior to experiments.

Fab fragments were cloned into pMT Puro (heavy chain) and pMT (light chain) insect cell expression vectors after the BiP signal sequence. The heavy chain Fab sequence was followed by a double StrepTagII tag for purification. Stable cell lines were produced by transfecting S2 cells in 6-well plates with heavy chain and light chain plasmids (1.6µg heavy chain and 0.4µg light chain DNA) using Effectene transfection reagent (Qiagen) according to the manufacturer’s protocol. Following transfection, cells were selected using Puromycin at 6 µg/mL and expanded into suspension cultures in Insect-XPRESS media. When cultures reached a density of 1*10^7^ cells/cm^2^, protein expression was induced by the addition of CuSO_4_ to a final concentration of 500µM. At 3-4 days after induction, supernatants were collected, adjusted to pH 8.0 with NaOH, and run over a StrepTactin Superflow affinity column (Cytiva) to purify Fab fragments. Fragments were eluted from the column in 2.5 mM d-desthiobiotin and buffer exchanged into PBS using a 10kDa molecular weight cutoff concentrator.

### Mouse immunization and antibody isolation

All animal experiments were approved by the Institutional Animal Care and Use Committee at the La Jolla Institute for Immunology (LJI) and strictly conducted according to the National Institutes of Health Guide for the Care and Use of Laboratory Animals. To produce murine antibodies, female BALB/cJ mice (Jackson labs, JAX:000651) were immunized with DNA plasmids encoding full-length PV-strain RABV-G in the pcDNA 3.1(-) vector under the CMV promoter. Endotoxin free DNA was produced using a Qiagen Endofree Gigaprep kit (#12391). Mice were injected intramuscularly in both quadriceps with 25 µg of 0.5 µg/mL plasmid DNA in PBS. Following injection of DNA, an AgilePulse Waveform Electroporator (BTX) was used to electroporate DNA for improved uptake. Mice were immunized with DNA on day 0 and again on day 14, and two mice were sacrificed on day 19 (5 days post-boost) to collect splenocytes for plasma cell isolation.

Spleens were briefly stored in DMEM with 5% FBS on ice prior to the start of the antibody discovery workflow. Spleens were homogenized via passage three times through a 40µm cell strainer (Corning, #431750) in DMEM with 5% FBS, pooled from both mice, and the resulting single cells were centrifuged at 1400rpm for 8 min at 4°C. Bulk splenocytes were subject to a non-plasma cell depletion cocktail using the CD138 + plasma cell isolation kit, (Miltenyi Biotec, #130-092-530) according to manufacturer’s instructions. Plasma cells were then purified using the EasySep Release Mouse CD138+ kit (STEMCELL Technologies, #100-0601) according to the manufacturer’s instructions. Following release of plasma cells from kit beads, cells were counted and loaded onto the Beacon Optifluidics System (Bruker Cellular Analysis) for rabies specific plasma cell isolation.

Biotinylated PV-strain RABV-G ectodomains were incubated with streptavidin coated polystyrene beads (Spherotech, # SVP5-60-5) and used to detect rabies-specific antibody secreting cells in nanoliter pens on OptoSelect 11kchips (Bruker Cellular Analysis). Secreted antibodies were detected over a 30 minute time course assay with a Goat anti Mouse IgG (H+L) AF Plus 594 secondary antibody (Invitrogen, # A32742) added to the antigen-coated beads resuspended in mouse plasma culture media (Bruker Cellular Analysis).

Thirteen RABV-G positive plasma cells were exported from the Beacon, three of which ultimately yielded unique, paired antibody heavy and light chain sequences encoding RABV-G positive antibodies. Cells were exported into 2x TCL buffer (Qiagen, #1070498) and mRNA was isolated using RNAClean XP Beads (Beckman Coulter, #A63987). Reverse transcription with Maxima H-(Thermo Scientific, # EP0753) was carried out according to the manufacturer’s protocol and using a template switch primer. Total cDNA was amplified using Platinum SuperFi II polymerase (Invitrogen, #12369050). After enzymatic cleanup, antibody heavy and light chain variable domains were amplified with two rounds of nested PCR using Platinum II Taq Hot-Start polymerase (Invitrogen, #14001014) using previously published primer sets(28). V_H_ and V_L_ domains were cloned into linearized human antibody expression vectors (human IgG1 and kappa light chain) using Gibson assembly (NEB, #E2621X) according to the manufacturer’s directions. Ligation reactions were transformed into 5-alpha F’I^q^ competent *E. coli* cells (NEB, #C2992I). DNA extraction was carried out with Qiagen buffer solutions and protocols. Plasmids were sequenced to ensure that the genes were in-frame, and that the cloned heavy and light chain variable domains matched PCR sequences.

Test expressions of each isolated antibody were carried out in 2.5 mL cultures of ExpiCHO cells cultured in 24-well blocks according to the manufacturer’s protocol for small-scale antibody expression. Supernatants containing secreted IgGs were collected 4-5 days post-transfection and were tested for RABV-G specificity via ELISA assay.

### RABV-G production and purification

Soluble PV-strain RABV-G ectodomains consisting of residues 1-439 were transiently expressed in adherent 293T cells, as previously described(19). Sequences were codon optimized for expression in human cell lines, and the first 19 amino acids of the sequence correspond to the glycoprotein’s native signal peptide. The glycoprotein ectodomain was either untagged (for cryo-EM complexes) or followed by a C-terminal double StrepTagII and Avi-tag, with linkers to add flexibility between tags (G-StrepTagII-GGGSGGGSGGGS-StrepTagII-GSGS-AviTag) (for all other experiments with soluble ectodomains). For both cases, the protein was cloned into a pcDNA 3.1(-) vector under the control of the CMV promoter. To produce glycoproteins, 293T cells were seeded into T75 flasks at a concentration of 4*10^4^ cells/cm^2^ and grown overnight in a standard, humidified CO_2_ incubator with 5% CO_2_ at 37°C. The following day, cells were transfected with 9.8µg plasmid per flask using Polyethyleneimine (PEI) (Polysciences) at a ratio of 3:1 DNA:PEI. Cell supernatant was collected and exchanged at two days post transfection, and collected again at four days post transfection.

Pooled cell supernatants were centrifuged to remove debris, adjusted to pH 8.0 with NaOH, and incubated with StrepTactin Superflow Plus (Qiagen) beads overnight on a shaker at 4°C. The following day, beads were collected in a gravity flow column, washed with PBS, and eluted in PBS with 25mM d-desthiobiotin. For bio-layer interferometry experiments and detection of rabies-specific plasma cells, glycoproteins were then concentrated to ∼4µM in a 100kDa molecular weight cutoff concentrator and biotinylated at the Avi Tag with BirA ligase (Avidity), according to the manufacturer’s instructions.

For cryo-EM structure determination, strep-tagged Fabs were bound to StrepTactin beads and washed to remove unbound antibodies. These beads were then incubated overnight with tissue culture supernatant containing untagged soluble RABV-G ectodomains. The following day, beads were collected in a gravity flow column, washed with PBS, and eluted with 25mM d-desthiobiotin. To trimerize glycoproteins or create high molecular weight complexes for cryo-EM, glycoprotein/antibody complexes were then incubated with additional Fab fragments in multiple (3-fold or higher) molar excess for one hour at room temperature. Following incubation, complexes were buffer exchanged into PBS and concentrated via three passes through a 500uL, 100 kDa MWCO concentrator.

### Cryo-EM specimen preparation and imaging

Complexes concentrated to ∼150µg/mL in PBS were mixed 1:3 with 0.36mM Lauryl maltose neopentyl glycol (LMNG) to a final concentration of 0.09mM immediately prior to freezing. 4µL of complexes were loaded onto C-flat 2/1 grids and frozen with a FEI Vitrobot at 100% humidity, a blot force of 0, and 10s blot time. Grids were imaged on a Titan Krios 300kV cryo-electron microscope with a Gatan K3 direct electron detector and energy filter, at a total dose of ∼50e^-^/Å^2^, 45,000x or 75,900x magnification and a pixel size of 1.1 or 0.66 (Table S1).

### Data processing

Cryo-EM maps were reconstructed in CryoSPARC(29) (Fig. S9 and S10). To avoid model bias, particles for each dataset were picked first with a blob picker, then with 2D classes generated within the dataset, and finally via Topaz 0.2.5a(30). Ab-initio 3D classes were also generated entirely within the dataset to avoid model bias. Maps were refined iteratively in CryoSPARC, using a combination of 2D and 3D classification (3D Hetero-refinement) to remove junk particles and debris, and 3D classification to account for particles with partial antibody occupancy on the RABV-G trimer. For maps except for the one with antibody A2, which had between 1-2 Fabs bound per trimer, trimers with partial antibody occupancy were excluded from the reconstruction. 3D homogenous refinement with C3 symmetry was used to reconstruct maps containing antibodies 4C12/4H3, 8C5, A4/7E8, and 10H5/7E8, with local refinement with C3 symmetry used to obtain the final map. For maps containing antibodies A2, A11, and RVC68, particles were symmetry expanded using C3 symmetry, then refined in 3D. The map for antibody A2 was refined using a combination of homogeneous refinement (C3), followed by C3 symmetry expansion, non-uniform refinement (C1), and local refinement (C1). The map for antibody A11 was refined first with homogenous refinement (C3), followed by C3 symmetry expansion, and local refinement (C1). The map for antibody RVC68 was refined as a monomer to account for flexibility in the fusion domain where RVC68 binds. It was refined first with a combination of homogenous, non-uniform, and local refinement (C3), then symmetry expanded (C3), and masked around the monomer/Fab complex, and refined with local refinement (C1) to obtain the final map. After reconstruction, maps were sharpened using DeepEMHancer (31), and models were built using COOT(32) and refined in Phenix(33). Maps and models were validated in Phenix and in the PDB(33) (Table S1).

### Neutralization Assays

Rabies pseudoviruses used in neutralization assays were prepared in 293T cells and titered on Vero cells. 293T cells were seeded in 6-well plates at a concentration of 9*10^5^ cells/well in 2 mL DMEM with 10% FBS (Day 1). Prior to seeding, wells were incubated with 0.1mg/mL poly-L-lysine (Sigma) for 1 hour at room temperature and washed with PBS. Cells were incubated overnight at 37°C and transfected with a plasmid encoding full-length PV-strain RABV-G the following day (Day 2). Each well was transfected with 1.5µg plasmid DNA and TransIT-LT1 (Mirus) at a 1:3 DNA:transfection reagent ratio, according to the manufacturer’s protocol. The following day (Day 3), transfected cells were infected with VSV-ΔG GFP parent virus (Kerafast) at an MOI of 1-2. Virus was incubated on cells in Opti-MEM (Gibco) with 2% FBS and 1x Penicillin/Streptomycin, in a total volume of 500µL/well for 1 hour, with plates rocked every 15 minutes. Following incubation, supernatant was removed and cells were washed once in Opti-MEM/2% FBS. 2mL of Opti-MEM/2% FBS were added to cells, and cells were incubated overnight at 37°C. At approximately 16 hours post-infection (Day 4), supernatant was collected from cells and frozen at -80°C.

Pseudoviruses were titered on Vero cells seeded in DMEM/10% FBS in 96-well plates at a concentration of 2*10^5^ cells/well. Plates were incubated at 37°C for at least 4 hours prior to infection to allow for cells to attach to plates. 1:10 serial dilutions of virus stocks were made by diluting thawed supernatant in Opti-MEM/2% FBS. Media was removed from cell plates and 50µL of serial dilutions were added to each well. Each serial dilution was plated in triplicate and plates were incubated overnight at 37°C. The following day, supernatant was removed from plates and cells were fixed in 4% paraformaldehyde/PBS and 20µg/mL Hoescht for 30 minutes at room temperature. Plates were washed twice in PBS to remove the fixative, and infected cells were counted in a ThermoFisher CX5 high-content cell imager to calculate viral titers.

For neutralization assays, Vero cells were seeded into 96-well plates, as described above. IgGs or Fab fragments were diluted to a starting concentration of 1-125µg/mL for IgGs and 0.24-30µg/mL for Fabs and serially diluted 1 part to 4 parts in Opti-MEM/2% FBS in 60µL in 96-well dilution plates. Following serial dilution, 15,000 ffu of pseudovirus in 60µL Opti-MEM/2% FBS was added to each dilution and plates were incubated at 37°C for 1 hour. After incubation, media was removed from Vero cells and replaced with 50µL of pseudovirus/antibody mixture.

Plates were incubated for 16 hours at 37°C, then cells were fixed and counted as described above to determine antibody neutralization titer. Three experimental replicates were performed, each consisting of two technical replicates.

### Bio-layer interferometry

Bio-layer interferometry experiments were performed on an Octet Red 384 instrument (Sartorius) using streptavidin biosensors (Sartorius). Biosensors were hydrated in kinetics buffer (PBS with 0.1% BSA and 0.02% CHAPS) for a minimum of 10 minutes prior to experiments.

Kinetics buffer was also used for all subsequent dilutions of proteins and washes in this protocol. For each experiment, the instrument was run in 8-biosensor mode, with 7 biosensors for data collection and one biosensor for reference subtraction. A baseline reading was collected for 30s, and then soluble RABV-G ectodomains biotinylated at the C-terminal Avi-tag were loaded onto biosensors to ∼1nm of binding over 5 minutes. Biosensors were washed for 1 minute, then incubated with Fab fragments at various concentrations (40, 20, 10, 5, 2.5, 1.25, and 0.625nM) in kinetics buffer for 5 minutes to measure association. Biosensors were then washed in kinetics buffer for 5 minutes to measure disassociation.

For competition assays, biosensors were loaded with RABV-G, as described above, and then saturated with 40nM of a Fab fragment. Biosensors were then washed in kinetics buffer and binding 40nM of additional Fab fragments from antibodies in our panel were measured.

Competition was determined by comparing Fab binding to RABV-G/Fab complexes with Fab binding to RABV-G alone.

Data was analyzed in the Octet Data Analysis 11.1 software using single reference subtraction, with biosensors with RABV-G and without Fabs serving as references for non-specific binding. Data was fit with a 1:1 binding kinetics model with global fitting, and kinetics experiments were performed in duplicate.

### Fusion Assays

For fusion assays, two populations of 293T cells each expressing part of a split mNeonGreen protein (mNG_1-10_ or mNG_11_) were co-seeded in 24-well plates in a 1:1 ratio and a concentration of 2*10^4^ cells/cm^2^ each (4*10^4^ cells/cm^2^ total). The following day, cells were transfected with full-length RABV-G using TransIT-LT1 transfection reagent at a 1:3 DNA:transfection reagent ratio, according to the manufacturer’s protocol. Three days post-transfection, cells were incubated with IgGs or Fabs (1.2-150 µg/mL IgG and 0.39-50 µg/mL Fab) in PBS for 1 hour at room temperature. Following incubation, cells were washed with PBS and incubated in citric acid buffer (0.1M citric acid/sodium citrate, pH 5.5) for 10 minutes at room temperature. Following incubation, citric acid buffer was removed from cells, and cells were washed in PBS and then incubated in DMEM with 10% FBS for 1 hour at 37°C to facilitate cell fusion. Cells were fixed in 4% paraformaldehyde/PBS for 30 minutes, washed with PBS, and imaged on a Keyence epifluorescence microscope to visualize syncytia formation.

### Binding and Uptake Assay

293T cells were seeded on 8-well glass bottom plates (IBIDI) at a density of 1*10^4^ cells/well and grown overnight at 37°C. The following day, rabies pseudoviruses 2*10^6^ ffu/well were incubated with IgG at a concentration of 4 times the IC90 for 1 hour on ice, then incubated with pre-cooled cells on coverslips for 1 hour on ice. Following incubation, cells were returned to 37°C for 15 minutes, then fixed in 4% paraformaldehyde/PBS for 15 minutes at room temperature and quenched in 20mM glycine/PBS for 5 minutes. Cells were permeabilized with 0.1% Triton X-100 in PBS for 5 minutes and blocked in 1% normal goat serum/PBS for 1 hour. Cells were then stained with stained with both 1µg/mL RVC68 and 1µg/mL RVA122 Fab fragments in PBS with 1% normal goat serum for 1 hour at room temperature, washed twice with PBS, and stained with 1:2,000 anti-human Fab DY549, 1:2,000 Phalloidin Alexa 488, and 20µg/mL Hoescht in PBS with 1% normal goat serum for 1 hour. Coverslips were then washed twice in PBS and mounted on slides with Prolong Antifade Gold mounting reagent (Invitrogen). Cells were imaged on a Zeiss 880 confocal microscope at 63x magnification.

### Immunofluorescence Assay

293T cells expressing half of a split mNeonGreen protein were seeded in 96-well plates at a concentration of 4*10^4^ cells/cm^2^ (2*10^4^ cells/cm^2^ each population) and incubated overnight at 37°C. The following day, cells were transfected with full-length PV-strain RABV-G or glycoprotein from Irkut lyssavirus, Ikoma lyssavirus, Eastern bat lyssavirus 1, Duvenhage lyssavirus, Mokola lyssavirus, or West Caucasian bat lyssavirus with TransIT-LT1 transfection reagent, as described above. Two days post-transfection, cells were either were fixed in 4% paraformaldehyde for 30 minutes for cell staining or incubated with 0.1M citric acid/sodium citrate buffer at pH 5.5 to induce fusion. For cell staining, cells were washed with PBS, and stained with either 5 µg/mL of one of the nine monoclonal antibodies from our panel, 1:500 HRIG, or 1:500 of a rabbit anti-RABV-G polyclonal antibody (a kind gift from Dr. Matthias Schnell, Thomas Jefferson University) in 1% BSA/PBS for 1 hour at room temperature. Cells were washed twice with PBS, then stained with 1:1,000 goat anti-human DY594 (Invitrogen) or goat anti-rabbit Alexa 568 and 20µg/mL Hoescht for 1 hour at room temperature.

## References

1. Organization TWH. 2026. Rabies. https://www.who.int/news-room/fact-sheets/detail/rabies. Accessed 01/02.

2. Zhang X, Zhu Z, Wang C. 2011. Persistence of rabies antibody 5 years after postexposure prophylaxis with vero cell antirabies vaccine and antibody response to a single booster dose. Clin Vaccine Immunol 18:1477–9.

3. De Benedictis P, Minola A, Rota Nodari E, Aiello R, Zecchin B, Salomoni A, Foglierini M, Agatic G, Vanzetta F, Lavenir R, Lepelletier A, Bentley E, Weiss R, Cattoli G, Capua I, Sallusto F, Wright E, Lanzavecchia A, Bourhy H, Corti D. 2016. Development of broad-spectrum human monoclonal antibodies for rabies post-exposure prophylaxis. EMBO Mol Med 8:407–21.

4. Hanna JN, Carney IK, Smith GA, Tannenberg AE, Deverill JE, Botha JA, Serafin IL, Harrower BJ, Fitzpatrick PF, Searle JW. 2000. Australian bat lyssavirus infection: a second human case, with a long incubation period. Med J Aust 172:597–9.

5. Fooks AR, McElhinney LM, Pounder DJ, Finnegan CJ, Mansfield K, Johnson N, Brookes SM, Parsons G, White K, McIntyre PG, Nathwani D. 2003. Case report: isolation of a European bat lyssavirus type 2a from a fatal human case of rabies encephalitis. J Med Virol 71:281–9.

6. Paweska JT, Blumberg LH, Liebenberg C, Hewlett RH, Grobbelaar AA, Leman PA, Croft JE, Nel LH, Nutt L, Swanepoel R. 2006. Fatal human infection with rabies-related Duvenhage virus, South Africa. Emerg Infect Dis 12:1965–7.

7. Familusi JB, Osunkoya BO, Moore DL, Kemp GE, Fabiyi A. 1972. A fatal human infection with Mokola virus. Am J Trop Med Hyg 21:959–63.

8. Roine RO, Hillbom M, Valle M, Haltia M, Ketonen L, Neuvonen E, Lumio J, Lahdevirta J. 1988. Fatal encephalitis caused by a bat-borne rabies-related virus. Clinical findings. Brain 111 ( Pt 6):1505–16.

9. Mire CE, Cross RW, Geisbert JB, Borisevich V, Agans KN, Deer DJ, Heinrich ML, Rowland MM, Goba A, Momoh M, Boisen ML, Grant DS, Fullah M, Khan SH, Fenton KA, Robinson JE, Branco LM, Garry RF, Geisbert TW. 2017. Human-monoclonal-antibody therapy protects nonhuman primates against advanced Lassa fever. Nat Med 23:1146–1149.

10. Mulangu S, Dodd LE, Davey RT, Jr., Tshiani Mbaya O, Proschan M, Mukadi D, Lusakibanza Manzo M, Nzolo D, Tshomba Oloma A, Ibanda A, Ali R, Coulibaly S, Levine AC, Grais R, Diaz J, Lane HC, Muyembe-Tamfum JJ, Group PW, Sivahera B, Camara M, Kojan R, Walker R, Dighero-Kemp B, Cao H, Mukumbayi P, Mbala-Kingebeni P, Ahuka S, Albert S, Bonnett T, Crozier I, Duvenhage M, Proffitt C, Teitelbaum M, Moench T, Aboulhab J, Barrett K, Cahill K, Cone K, Eckes R, Hensley L, Herpin B, Higgs E, Ledgerwood J, Pierson J, Smolskis M, Sow Y, Tierney J, Sivapalasingam S, Holman W, Gettinger N, et al. 2019. A Randomized, Controlled Trial of Ebola Virus Disease Therapeutics. N Engl J Med 381:2293–2303.

11. Cross RW, Hastie KM, Mire CE, Robinson JE, Geisbert TW, Branco LM, Ollmann Saphire E, Garry RF. 2019. Antibody therapy for Lassa fever. Curr Opin Virol 37:97–104.

12. Zorzan M, Castellan M, Gasparotto M, Dias de Melo G, Zecchin B, Leopardi S, Chen A, Rosato A, Angelini A, Bourhy H, Corti D, Cendron L, De Benedictis P. 2023. Antiviral mechanisms of two broad-spectrum monoclonal antibodies for rabies prophylaxis and therapy. Front Immunol 14:1186063.

13. Gaudin Y, Raux H, Flamand A, Ruigrok RW. 1996. Identification of amino acids controlling the low-pH-induced conformational change of rabies virus glycoprotein. J Virol 70:7371–8.

14. Gaudin Y, Ruigrok RW, Knossow M, Flamand A. 1993. Low-pH conformational changes of rabies virus glycoprotein and their role in membrane fusion. J Virol 67:1365–72.

15. Gaudin Y, Tuffereau C, Durrer P, Flamand A, Ruigrok RW. 1995. Biological function of the low-pH, fusion-inactive conformation of rabies virus glycoprotein (G): G is transported in a fusion-inactive state-like conformation. J Virol 69:5528–34.

16. Roche S, Gaudin Y. 2002. Characterization of the equilibrium between the native and fusion-inactive conformation of rabies virus glycoprotein indicates that the fusion complex is made of several trimers. Virology 297:128–35.

17. Raux H, Coulon P, Lafay F, Flamand A. 1995. Monoclonal antibodies which recognize the acidic configuration of the rabies glycoprotein at the surface of the virion can be neutralizing. Virology 210:400–8.

18. Yang F, Lin S, Ye F, Yang J, Qi J, Chen Z, Lin X, Wang J, Yue D, Cheng Y, Chen Z, Chen H, You Y, Zhang Z, Yang Y, Yang M, Sun H, Li Y, Cao Y, Yang S, Wei Y, Gao GF, Lu G. 2020. Structural Analysis of Rabies Virus Glycoprotein Reveals pH-Dependent Conformational Changes and Interactions with a Neutralizing Antibody. Cell Host Microbe 27:441–453 e7.

19. Callaway HM, Zyla D, Larrous F, de Melo GD, Hastie KM, Avalos RD, Agarwal A, Corti D, Bourhy H, Saphire EO. 2022. Structure of the rabies virus glycoprotein trimer bound to a prefusion-specific neutralizing antibody. Sci Adv 8:eabp9151.

20. Evans JS, Horton DL, Easton AJ, Fooks AR, Banyard AC. 2012. Rabies virus vaccines: is there a need for a pan-lyssavirus vaccine? Vaccine 30:7447–54.

21. Hellert J, Buchrieser J, Larrous F, Minola A, de Melo GD, Soriaga L, England P, Haouz A, Telenti A, Schwartz O, Corti D, Bourhy H, Rey FA. 2020. Structure of the prefusion-locking broadly neutralizing antibody RVC20 bound to the rabies virus glycoprotein. Nat Commun 11:596.

22. Ng WM, Fedosyuk S, English S, Augusto G, Berg A, Thorley L, Haselon AS, Segireddy RR, Bowden TA, Douglas AD. 2022. Structure of trimeric pre-fusion rabies virus glycoprotein in complex with two protective antibodies. Cell Host Microbe 30:1219–1230 e7.

23. Cao L, Qu J, Zhang C, Chen R, Wang J, Fehlner-Gardiner C, Niezgoda M, Satheshkumar PS, Wang X, Tsao E. 2025. Structural basis for broad neutralization of rabies virus by an antibody cocktail SYN023. Emerg Microbes Infect 14:2547724.

24. Dessain SK. 2018. Compositions comprising antibodies to rabies virus and the uses thereofWO 2019/227039 A1.

25. Davide Corti HB. 2015. Antibodies that potently neutralize rabies virus and other lyssaviruses and uses thereof. United States.

26. Aditham AK, Radford CE, Carr CR, Jasti N, King NP, Bloom JD. 2025. Deep mutational scanning of rabies glycoprotein defines mutational constraint and antibody-escape mutations. Cell Host Microbe 33:988–1003 e10.

27. de Melo GD, Sonthonnax F, Lepousez G, Jouvion G, Minola A, Zatta F, Larrous F, Kergoat L, Mazo C, Moigneu C, Aiello R, Salomoni A, Brisebard E, De Benedictis P, Corti D, Bourhy H. 2020. A combination of two human monoclonal antibodies cures symptomatic rabies. EMBO Mol Med 12:e12628.

28. Tiller T, Busse CE, Wardemann H. 2009. Cloning and expression of murine Ig genes from single B cells. J Immunol Methods 350:183–93.

29. Punjani A, Rubinstein JL, Fleet DJ, Brubaker MA. 2017. cryoSPARC: algorithms for rapid unsupervised cryo-EM structure determination. Nat Methods 14:290–296.

30. Bepler T, Morin A, Rapp M, Brasch J, Shapiro L, Noble AJ, Berger B. 2019. Positive-unlabeled convolutional neural networks for particle picking in cryo-electron micrographs. Nat Methods 16:1153–1160.

31. Sanchez-Garcia R, Gomez-Blanco J, Cuervo A, Carazo JM, Sorzano COS, Vargas J. 2021. DeepEMhancer: a deep learning solution for cryo-EM volume post-processing. Commun Biol 4:874.

32. Emsley P, Lohkamp B, Scott WG, Cowtan K. 2010. Features and development of Coot. Acta Crystallogr D Biol Crystallogr 66:486–501.

33. Williams CJ, Headd JJ, Moriarty NW, Prisant MG, Videau LL, Deis LN, Verma V, Keedy DA, Hintze BJ, Chen VB, Jain S, Lewis SM, Arendall WB, 3rd, Snoeyink J, Adams PD, Lovell SC, Richardson JS, Richardson DC. 2018. MolProbity: More and better reference data for improved all-atom structure validation. Protein Sci 27:293–315.

