## Supplemental Figures and Tables for "Antigenic landscape of rabies and related lyssaviruses revealed by cryo-EM"

### Supplemental Information

A

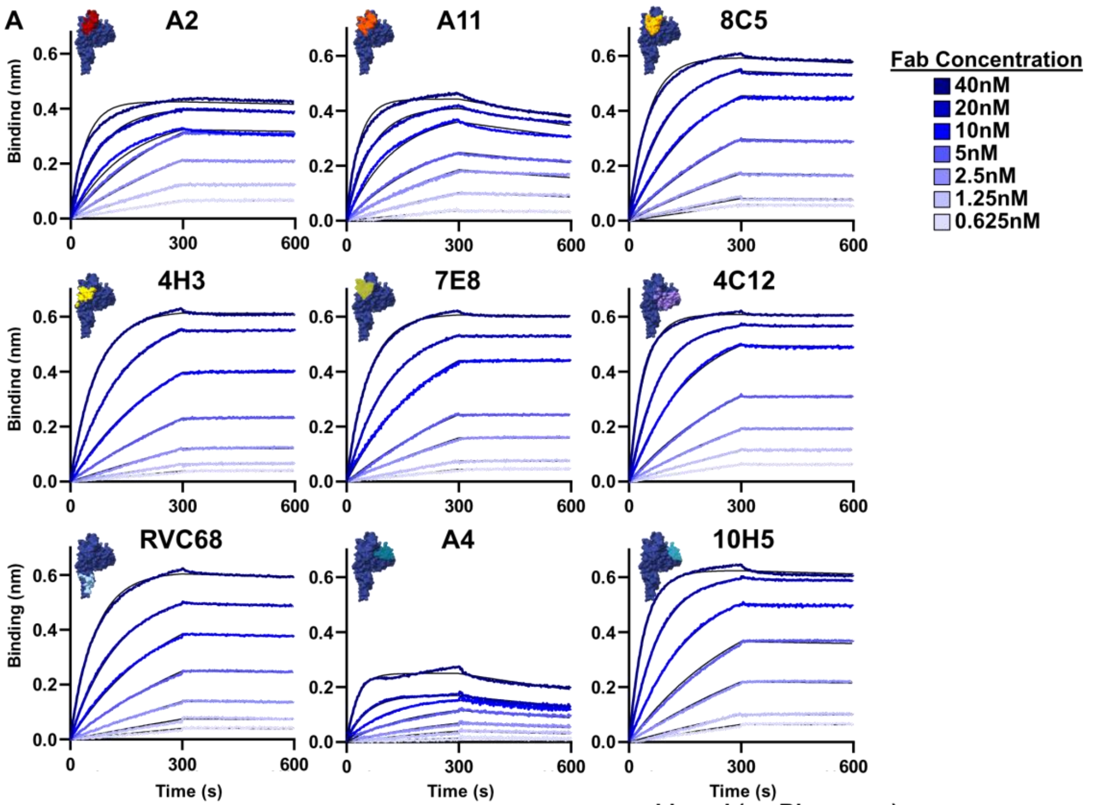


B

**Binding Kinetics**

| | KD (M) | Ka (1/M*s) | $K_d$ (1/s) |
|---|---|---|---|
| **A2** | 1.13E-10 | 4.89E+05 | 5.52E-05 |
| **A11** | 8.01E-10 | 7.49E+05 | 6.00E-04 |
| **8C5** | 6.21E-11 | 6.80E+05 | 4.22E-05 |
| **4H3** | 1.33E-10 | 6.25E+05 | 8.30E-05 |
| **7E8** | 2.78E-10 | 6.79E+05 | 2.00E-04 |
| **4C12** | 2.82E-10 | 8.49E+04 | 2.00E-04 |
| **RVC68** | 1.52E-10 | 3.81E+05 | 5.79E-05 |
| **A4** | 1.54E-09 | 8.51E+05 | 1.30E-03 |
| **10H5** | 9.81E-11 | 8.52E+05 | 8.30E-05 |

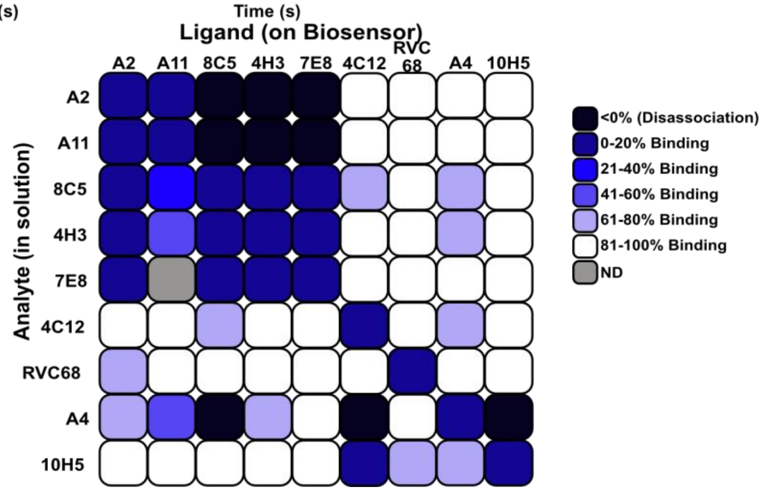


**Supplemental Figure 1. Antibodies bind RABV-G with nanomolar or higher affinity.** Binding kinetics of Fab fragments to RABV-G, measured by bio-layer interferometry. Representative binding curves (A) and binding kinetics (B) are shown. Competition between Fab fragments from different antibodies (C), indicates that antibodies with overlapping binding footprints inhibit each other's binding or cause antibodies to disassociate from RABV-G.

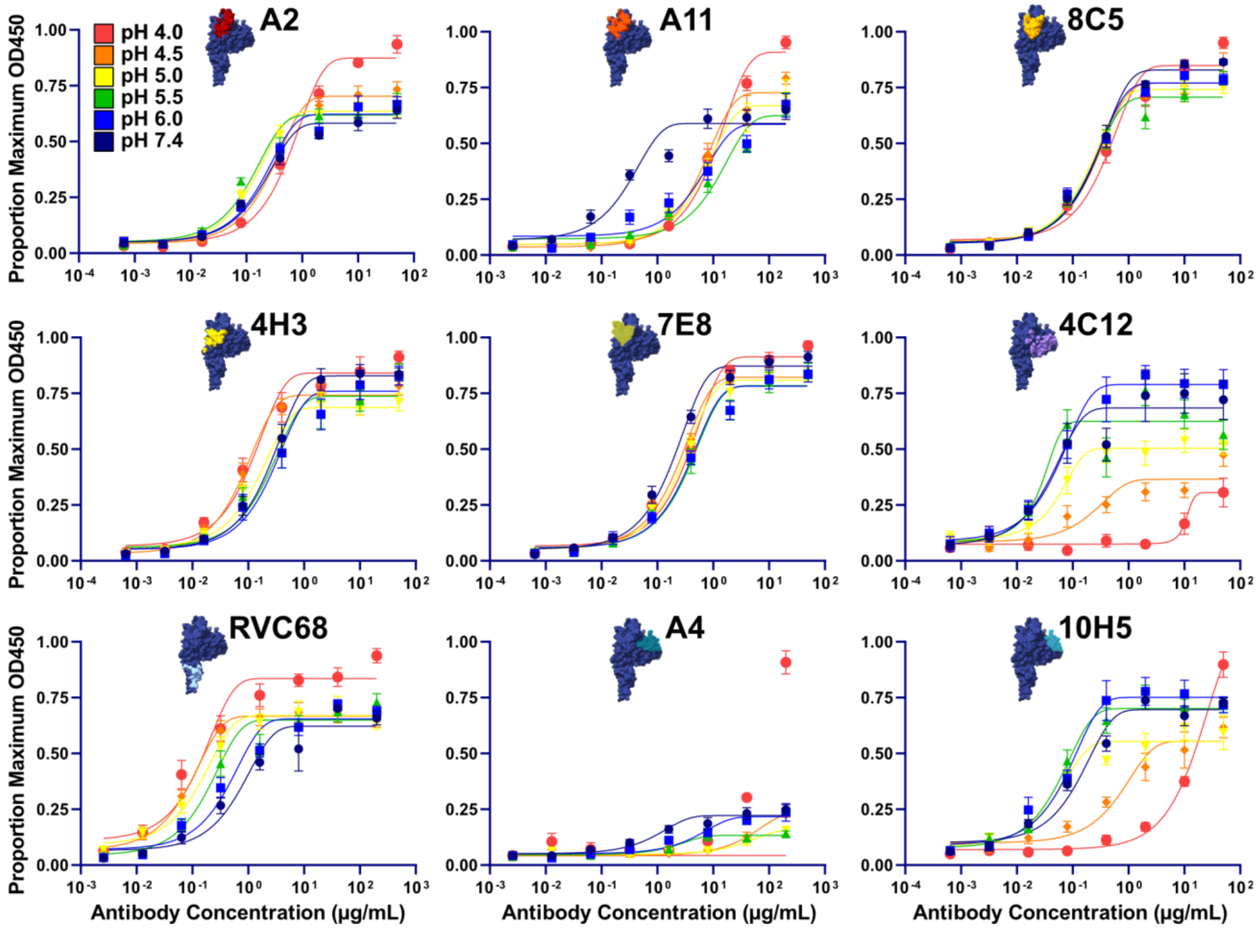


10 **Supplemental Figure 2. Site III mAbs are pH sensitive, but site II/IV mAbs are not.** ELISA assay showing binding of IgGs to RABV-G over a range of pHs. Binding for each antibody was normalized to the maximum OD450 seen for that antibody across all pH values and three experimental replicates.

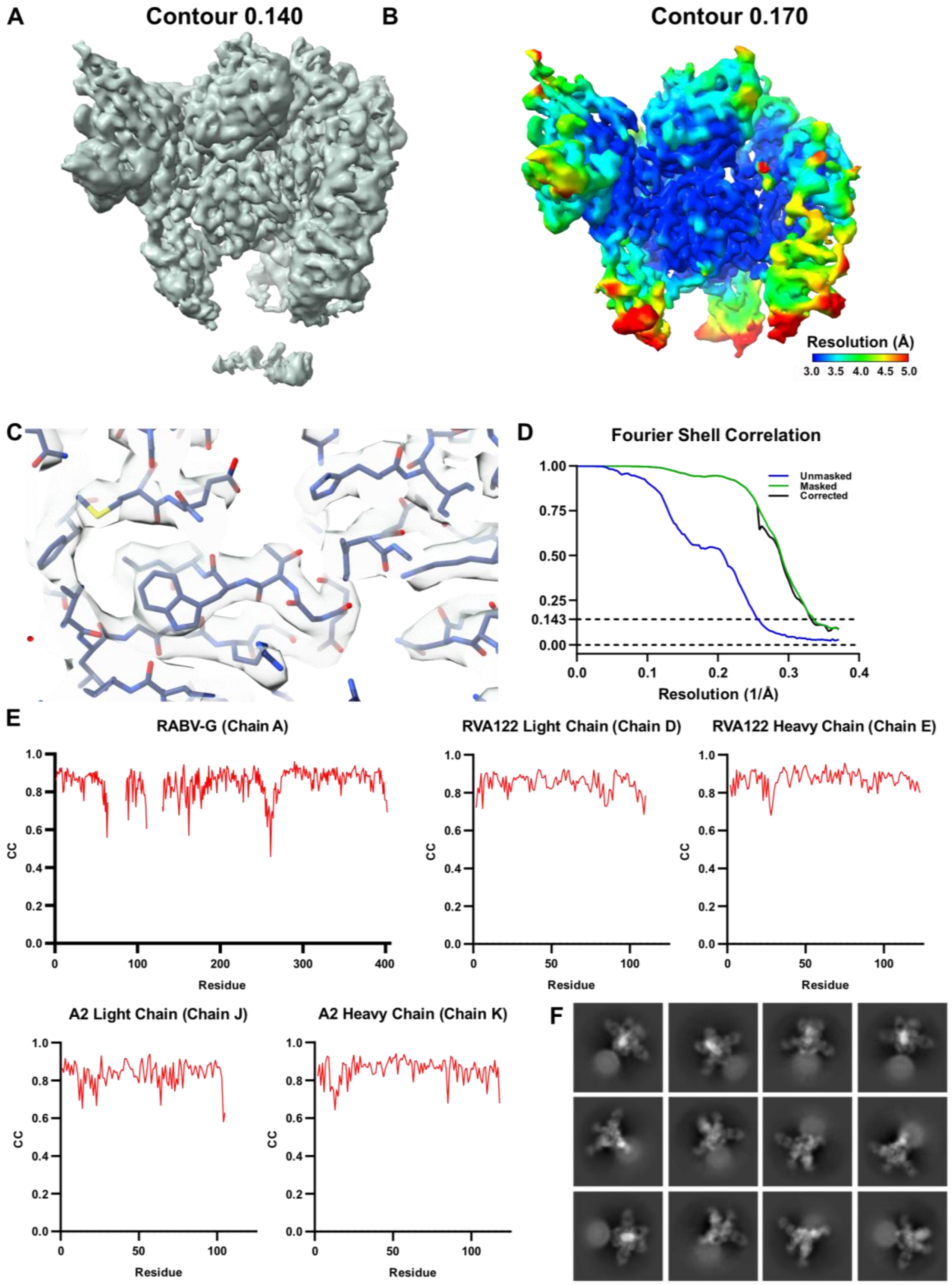


**Supplemental Figure 3. Map and model statistics for mAb A2.** Cryo-EM map at high contour (A), and low contour (B) with color-coding for local resolution are shown. Atomic model docked into density (C), FSC curves (D), and Phenix map/model correlation coefficient graphs for unique chains (E) are also shown. Site III binding monoclonal antibody RVA122 was also included in complexes to stabilize RABV-G pre-fusion trimers for high-resolution imaging. Representative 2D classes are shown in (F).

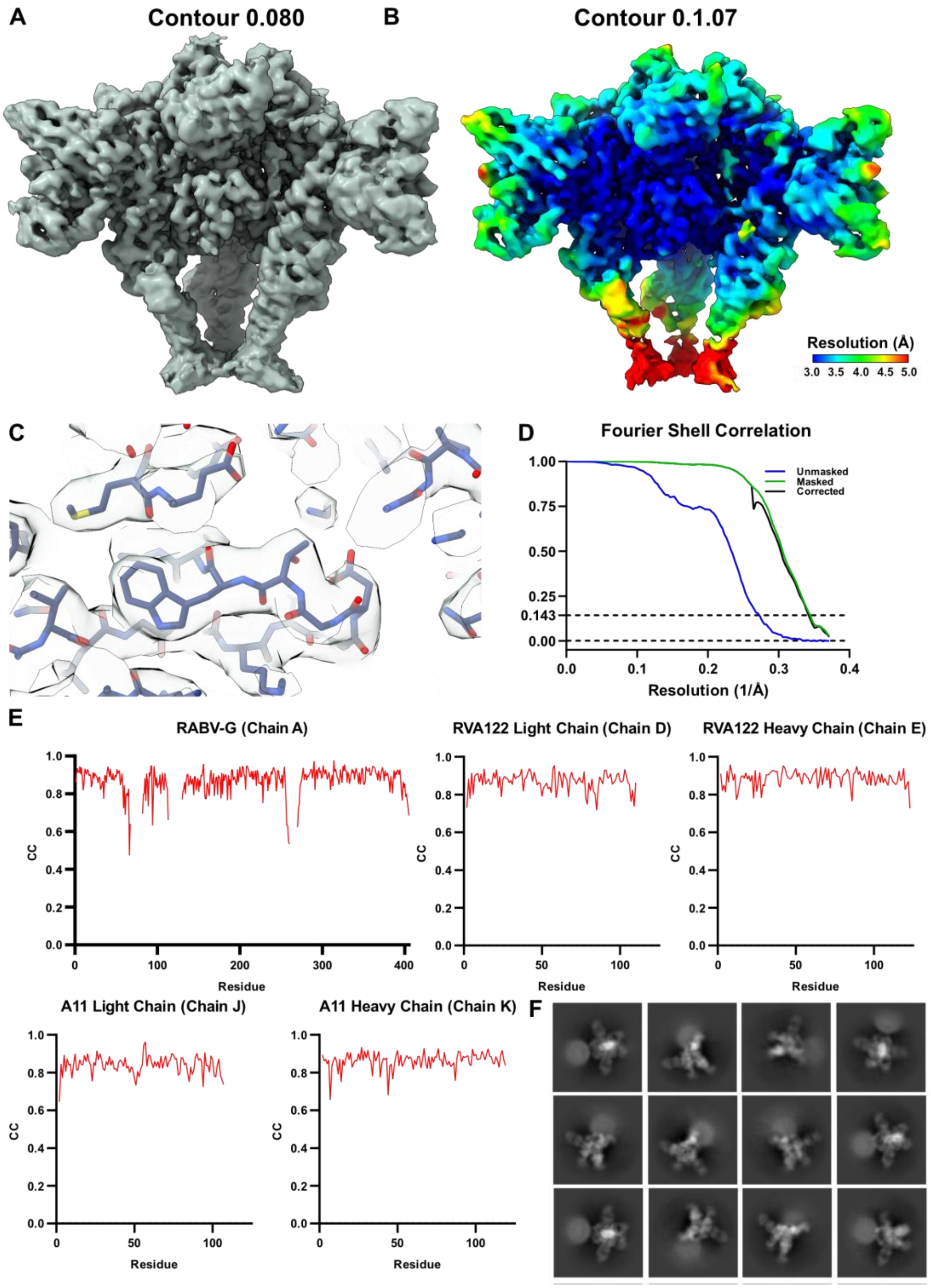


**Supplemental Figure 4. Map and model statistics for mAb A11.** Cryo-EM map at high contour (A), and low contour (B) with color-coding for local resolution are shown. Atomic model docked into density (C), FSC curves (D), and Phenix map/model correlation coefficient graphs for unique chains (E) are also shown. Site III binding monoclonal antibody RVA122 was also included in complexes to stabilize RABV-G pre-fusion trimers for high-resolution imaging. Representative 2D classes are shown in (F).

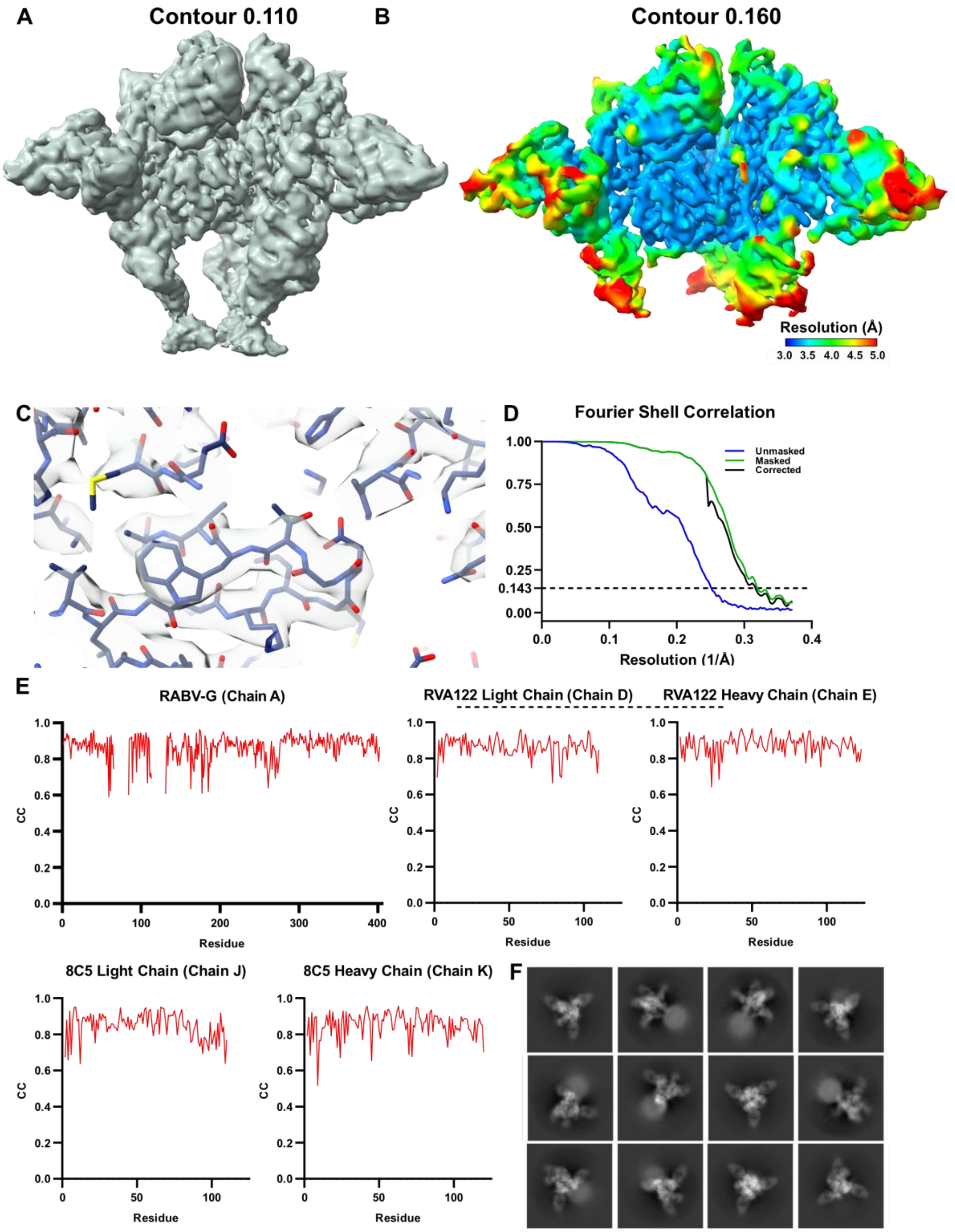


**Supplemental Figure 5. Map and model statistics for mAb 8C5.** Cryo-EM map at high contour (A), and low contour (B) with color-coding for local resolution are shown. Atomic model docked into density (C), FSC curves (D), and Phenix map/model correlation coefficient graphs for unique chains (E) are also shown. Site III binding monoclonal antibody RVA122 was also included in complexes to stabilize RABV-G pre-fusion trimers for high-resolution imaging. Representative 2D classes are shown in (F).

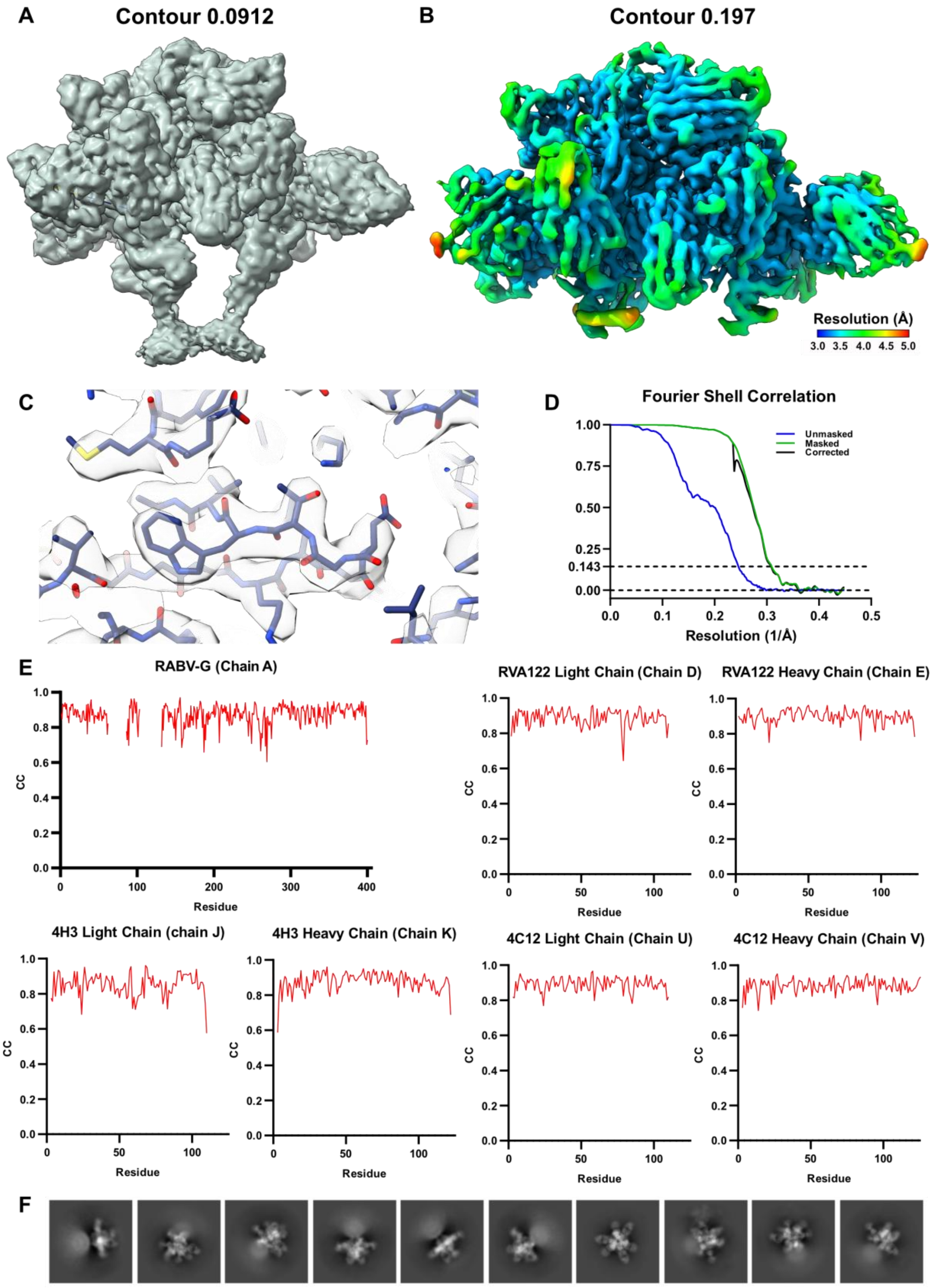


**Supplemental Figure 6. Map and model statistics for mAbs 4C12 and 4H3.** Cryo-EM map at high contour (A), and low contour (B) with color-coding for local resolution are shown. Atomic model docked into density (C), FSC curves (D), and Phenix map/model correlation coefficient graphs for unique chains (E) are also shown. Site III binding monoclonal antibody RVA122 was also included in complexes to stabilize RABV-G pre-fusion trimers for high-resolution imaging. Representative 2D classes are shown in (F).

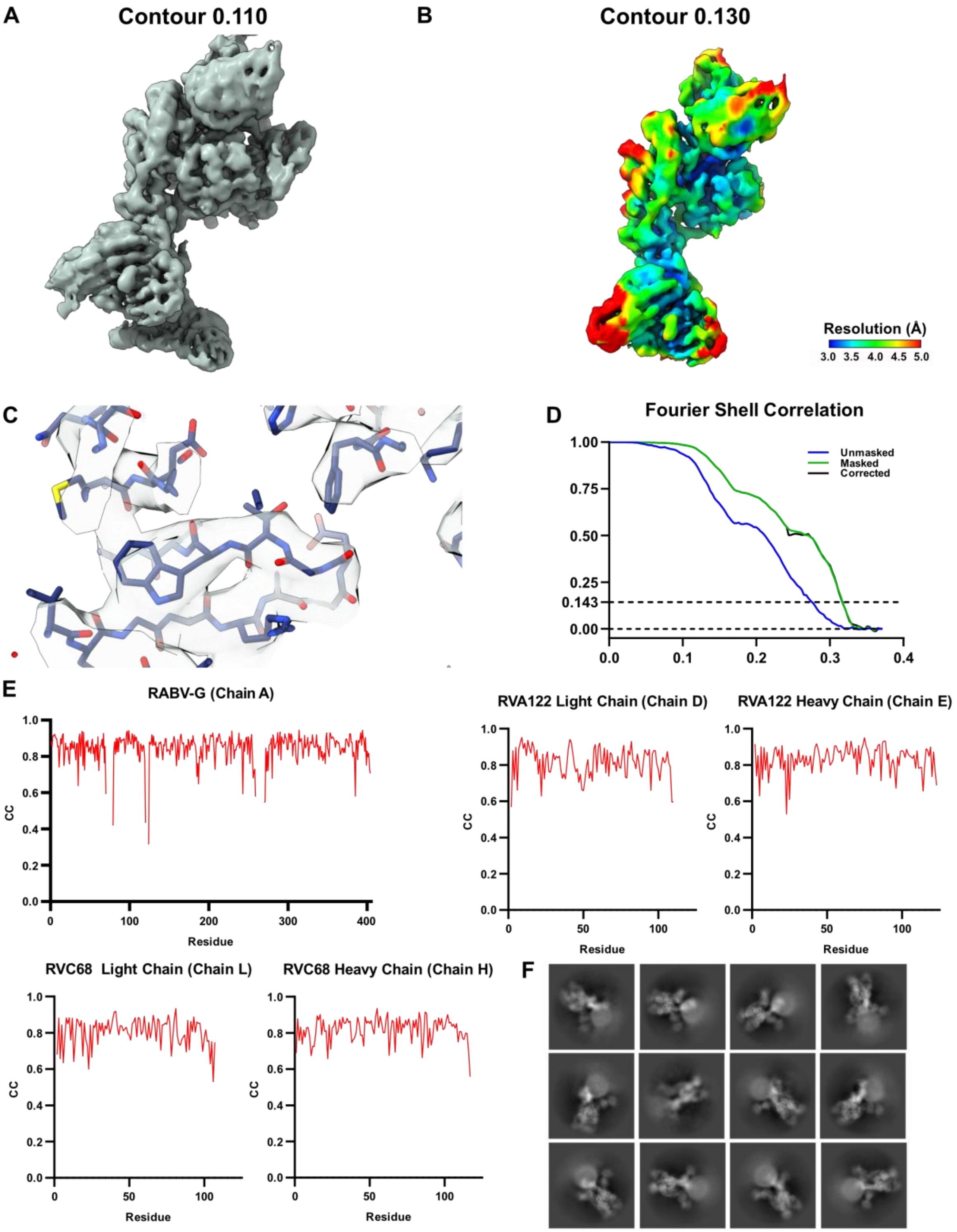


**Supplemental Figure 7. Map and model statistics for mAb RVC68.** Cryo-EM map at high contour (A), and low contour (B) with color-coding for local resolution are shown. Atomic model docked into density (C), FSC curves (D), and Phenix map/model correlation coefficient graphs for unique chains (E) are also shown. While this complex formed stable trimers, flexibility in the fusion domain where RVC68 binds necessitated processing the glycoprotein as a monomer with extensive 3D classification sorting to solve a high-resolution structure. Site III binding monoclonal antibody RVA122 was also included in this complex. Representative 2D classes are shown in (F).

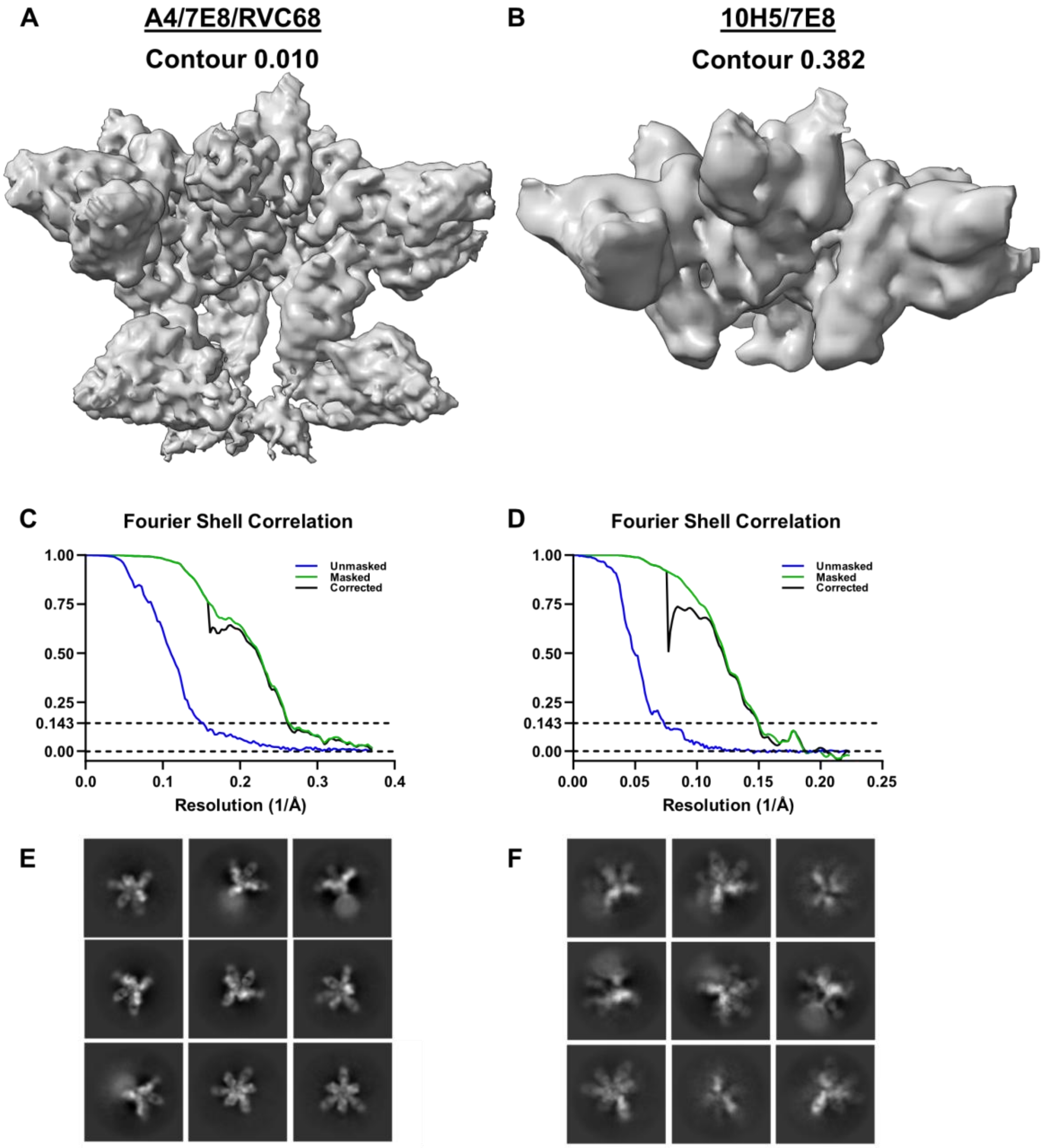


**Supplemental Figure 8. Map statistics for low-resolution maps of mAbs A4, 7E8, and 10H5.** Cryo-EM maps at high contour (A and C), and FSC curves (B and D) for mAbs A4, 7E8, and 10H5 are shown. Complexes consist of RABV-G trimers with either antibodies A4, 7E8, and RVC68 or antibodies 10H5 and 7E8. Representative 2D classes are shown in (E) and (F).

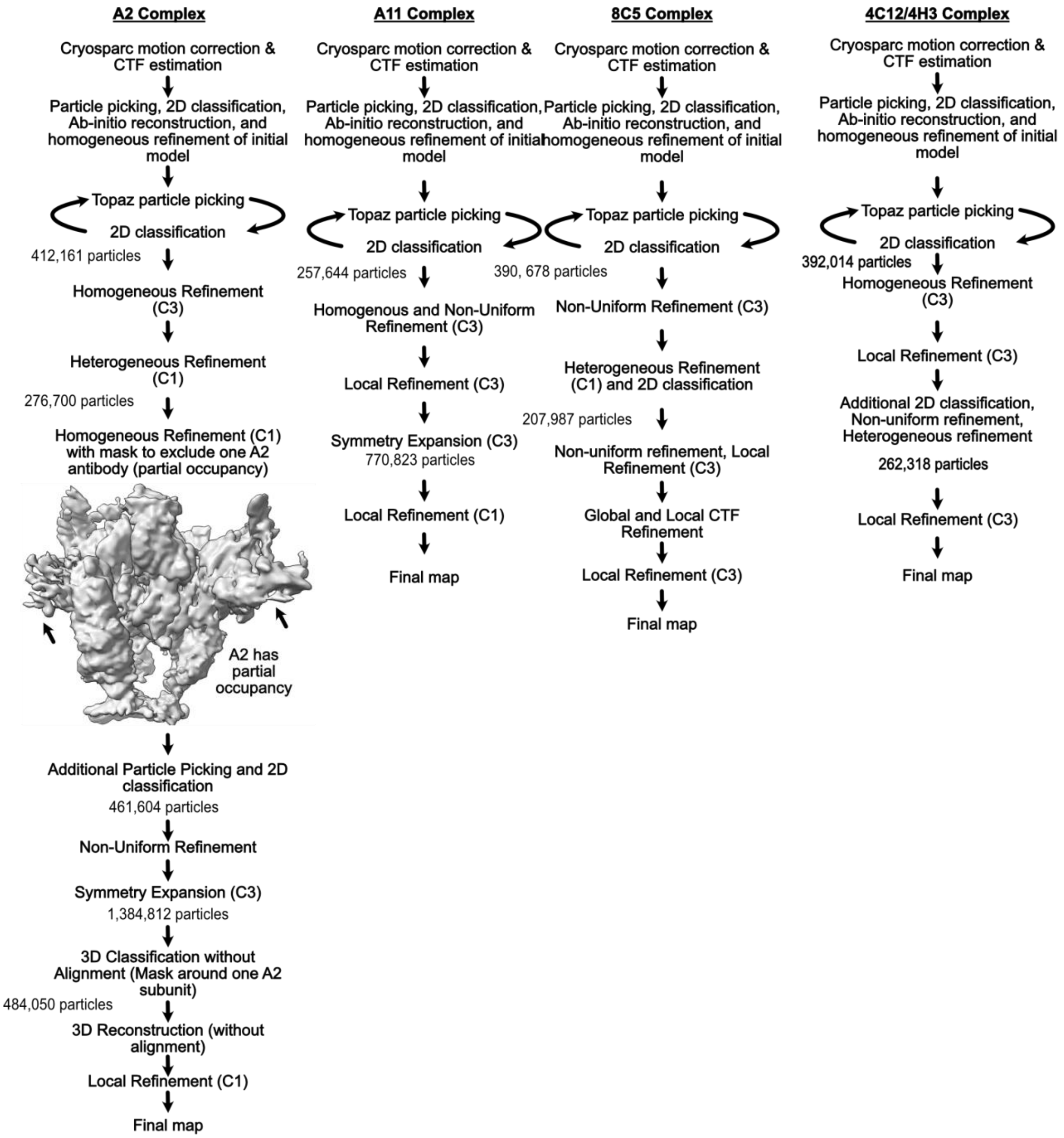


**Supplemental Figure 9. 3D Reconstruction workflow for cryo-EM maps.** Workflows for 3D map reconstruction for RABV-G in complex with antibodies A2, A11, 8C5, and 4C12/4H3.

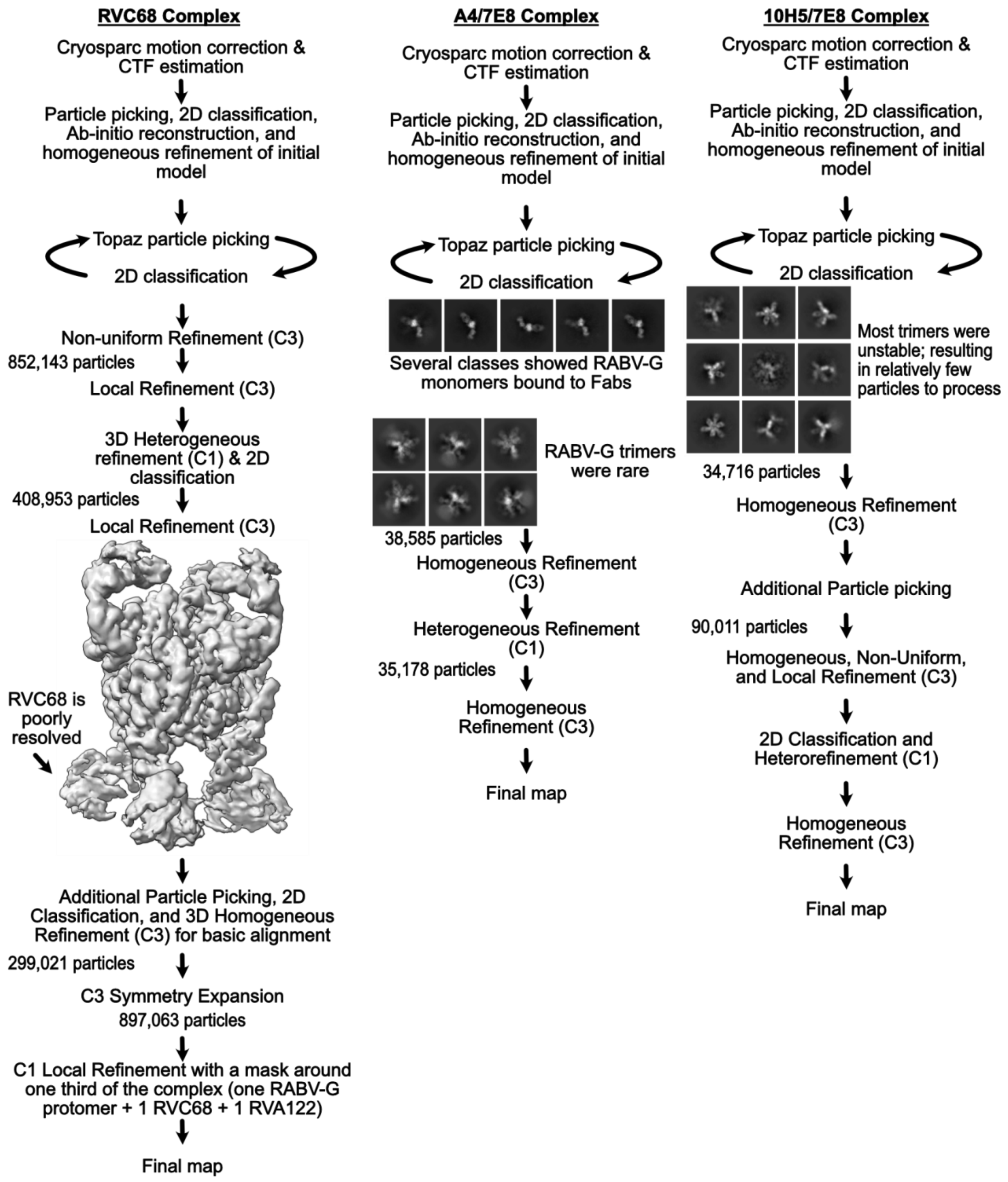


65

**Supplemental Figure 10. 3D reconstruction workflow for cryo-EM maps.** Workflows for 3D map reconstruction of RABV-G in complex with antibodies RVC68, A4/7E8, and 10H5/7E8.

| **Map Statistics** | **4C12 and 4H3 with RABV-G** | **A2 with RABV-G** | **A11 with RABV-G** | **8C5 with RABV-G** | **RVC68 with RABV-G** | **A4 and 7E8 with RABV-G** | **10H5 and 7E8 with RABV-G** |
|---|---|---|---|---|---|---|---|
| Movies | 4,167 | 11,395 | 15,077 | 10,530 | 8,416 | 12,794 | 17,703 |
| Particles | 262,318 | 484,050 | 770,823 | 208,987 | 897,063 | 35,178 | 25,156 |
| B-factor | 111.4 | 96.2 | 107.3 | 112.3 | 87.2 | 50.6 | 385.8 |
| Symmetry | C3 | C1 (C3 symmetry expanded) | C1 (C3 symmetry expanded) | C3 | C1 (C3 symmetry expanded) | C3 | C3 |
| Resolution (Å) (0.143 threshold) | 3.24 | 3.01 | 2.92 | 3.16 | 3.15 | 3.83 | 6.69 |
| Box size (px) | 400 | 512 | 512 | 512 | 512 | 512 | 620 |
| Pixel size (Å/px) | 1.1 | 0.66 | 0.66 | 0.66 | 0.66 | 0.66 | 1.1 |
| Microscope | Titan Krios | Titan Krios | Titan Krios | Titan Krios | Titan Krios | Titan Krios | Titan Krios |
| Detector | Gatan K3 | Gatan K3 | Gatan K3 | Gatan K3 | Gatan K3 | Gatan K3 | Gatan K3 |
| Total dose $(e^-/Å^2)$ | 50 | 50 | 50 | 50 | 50 | 50 | 50 |
| Frames | 50 | 50 | 50 | 50 | 50 | 25 | 25 |
| Dose per frame $(e^-/Å^2)$ | 1 | 1 | 1 | 1 | 1 | 2 | 2 |

| **Molprobity statistics** | | | | | | | |
|---|---|---|---|---|---|---|---|
| All-atom clash score | 5.17 | 4.87 | 3.67 | 5.3 | 4.75 | N/A | N/A |
| Ramachandran plot: | | | | | | N/A | N/A |
| Outliers | 0.00% | 0.00% | 0.00% | 0.00% | 0 | N/A | N/A |
| Allowed | 1.73% | 2.23% | 1.62% | 3.04% | 2.32% | N/A | N/A |
| Favored | 98.27% | 97.77% | 98.38% | 96.96% | 97.68% | N/A | N/A |
| C-beta outliers | 0.00% | 0.00% | 0.00% | 0.00% | 0 | N/A | N/A |
| Rotamer outliers | 0.00% | 0.06% | 0.00% | 0.00% | 0 | N/A | N/A |
| Peptide plane: | | | | | | N/A | N/A |
| Cis-proline | 0.00% | 0.00% | 0.00% | 0.00% | 2.2 | N/A | N/A |
| Cis-general | 0.00% | 0.00% | 0.00% | 0.00% | 0 | N/A | N/A |
| Twisted proline | 0.00% | 0.00% | 0.00% | 0.00% | 0 | N/A | N/A |
| Twisted general | 0.00% | 0.00% | 0.00% | 0.00% | 0 | N/A | N/A |

**Table S1. Cryo-EM map and model statistics.**
